# The parietal operculum preferentially encodes heat pain and not salience

**DOI:** 10.1101/581504

**Authors:** Björn Horing, Christian Sprenger, Christian Büchel

**Affiliations:** Affective Neuroscience Group, Department of Systems Neuroscience, University Medical Center Hamburg-Eppendorf

## Abstract

Substantial controversy exists as to which part of brain activity is genuinely attributable to pain-related percepts, and which activity is due to general aspects of sensory stimulation, such as its salience, or the accompanying arousal. The challenge posed by this question rests largely in the fact that pain per se exhibits highly intense but unspecific characteristics. These therefore should be matched by potential control conditions. Here, we used a unique combination of functional magnetic resonance imaging, behavioral and autonomic measures to address this longstanding debate in pain research.

Subjects rated perceived intensity in a sequence alternating between heat and sound stimuli. Neuronal activity was monitored using fMRI. Either modality was presented in six different intensities, three of which lay above the pain threshold (for heat) or the unpleasantness threshold (for sound). We performed our analysis on 26 volunteers in which psychophysiological responses (as per skin conductance responses) did not differ between the two stimulus modalities. Having thus ascertained a comparable amount of stimulation-related but unspecific activation, we analyzed stimulus response functions after painful stimulation, and contrasted them with those of the matched acoustic control condition. Furthermore, analysis of fMRI data was performed on the brain surface to circumvent blurring issues stemming from the close proximity of several regions of interest located in heavily folded cortical areas. We focused our analyses on insular and periinsular regions which are strongly involved in processing of painful stimuli. We employed an axiomatic approach to determine areas showing higher activation in painful compared to non-painful heat, and at the same time showing a steeper stimulus response function for painful heat as compared to unpleasant sound. Intriguingly, an area in the posterior parietal operculum emerged whose response showed a pain preference after satisfying all axiomatic constraints.

This result has important implications for the interpretation of functional imaging findings in pain research, as it clearly demonstrates that there are areas whose activity following painful stimulation is not due to general attributes or results of sensory stimulation, such as salience or arousal. Conversely, several areas did not conform to the formulated axioms to rule out general factors as explanations.

## Introduction

Pain is a multidimensional experience, including sensory-discriminative, affective-motivational, cognitive-evaluative as well as motor components [1], and is defined as “an unpleasant sensory and emotional experience associated with actual or potential tissue damage, or described in terms of such damage” [2]. Following the advent of brain imaging, recurring patterns of brain activity following painful stimuli were identified, comprising primary and secondary somatosensory cortices, cingulate cortices, as well as the insular subregions, among other structures [3, 4].

This activity has frequently been attributed to pain per se. However, precisely because pain is a composite sensation, the utility of assigning it a fixed set of brain areas as a simplistic “pain matrix” has been contested [3, 4]. The notion that some of the observed activation may or may not be exclusively pain-related has eventually led to a direct challenge to the concept of such a matrix [5, 6]. These studies have provided evidence that in many cortical regions, activation is observed following both painful and non-painful (such as tactile or auditory) stimuli. In fact, general processes have long been posited as alternative explanations – for example, stimulus anticipation [7], magnitude estimation [8], or stimulus salience [5]. These contributions have led to lively controversy [9–12], and a large number of studies continues to address related issues using multiple modalities, and both univariate and multivariate approaches [13–18]. The status of the challenge to the “pain matrix” concept has recently been revisited [19]. This review reemphasized that great care should be taken experimentally to match non-painful control modalities, which has frequently been neglected in previous studies.

In addition to the question of general, modality-independent stimulus characteristics, many experiments have relied on the use of single stimulus intensities to characterize neuronal responses, when using painful stimulation and compared these responses to a non-painful control condition. However, such approaches disregard the possibility of modality-specific baseline activation, further compounding the issue to properly account for nonspecific activation [12]. A possible solution is to employ multiple stimulus intensities, which allows for the characterization of modality-specific stimulus-response functions [20–22], and a comparison of these between modalities.

Here, we address these issues, and present a novel approach that allows to directly test whether there are cortical regions that can be viewed as “salience detectors”, or show preferential pain processing. Please note that the term “salience” here and in the following does not refer to any narrowly defined physiological or cognitive construct, but is instead meant as generic term referencing the unspecific, modality-independent results of sensory stimulation. We employed heat and sound as stimulus modalities. Stimuli were presented in alternating modalities in a within-subjects design. Of each modality, we used six graded intensities – three below and three above the pain and unpleasantness threshold, respectively. This allowed us to determine stimulus-response functions of physical intensities or their percepts, and relate those to neuronal activity [20–23]. Importantly, auditory and thermal intensities were calibrated to match using an objective autonomic measure (skin conductance responses; SCR) [24].

We paid particular attention to insular and periinsular regions, especially the posterior insula and the parietal operculum (the secondary somatosensory cortex), all of which have been reported as cortical areas that are active early after painful stimulation [4,11,25–27].

To define areas as preferentially pain-processing, our analyses followed an axiomatic approach which posits several logical conditions to be met to make a valid inference (see [28], for a similar approach in pain avoidance). Within this rigorous approach, we formulated the following set of conditions to preclude the possibility that activity in an area could be explained by salience alone: The effect of painful stimulation should be larger than that of non-painful heat (axiom 1); the effect of painful stimulation should be larger than that of (salience-matched) unpleasant sound (axiom 2); the relationship of ratings and BOLD should be stronger for painful heat than for non-painful heat (axiom 3); the positive relationship of pain ratings and BOLD responses should be stronger for painful heat than for (salience-matched) unpleasant sound (axiom 4).

## Results

Heat stimuli were presented using a CHEPS thermode, sounds were 1kHz beeps presented binaurally via headphones. For a brief overview, see Fig 1. Details are provided in the Materials and Methods section.

**Fig 1.**
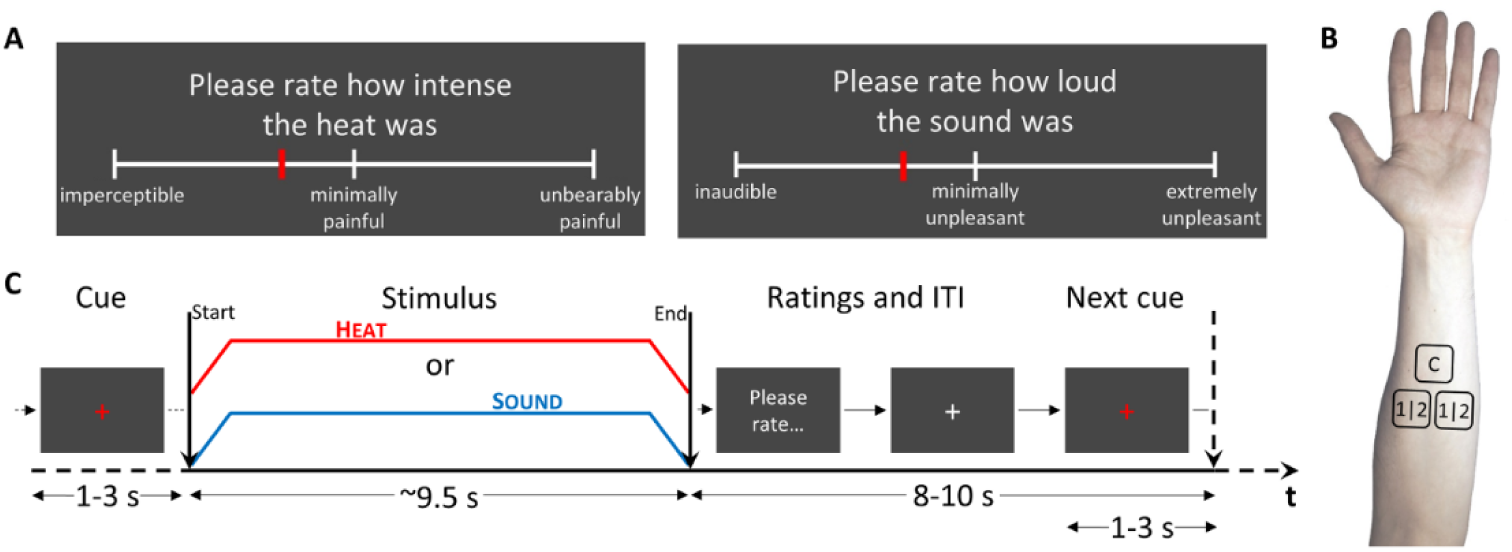
Experimental procedures. **A.** Visual analogue scales (VAS) for rating heat (left) and sound stimuli (right). The midpoint signifies the pain threshold, corresponding to a score of 0 in a conventional VAS. **B.** Thermode arrangement on a subject’s forearm. Patch C was used for calibration, patches 1|2 were used in counterbalanced fashion for experimental sessions 1 and 2. **C.** Protocol by time. A visual cue (white fixation cross turning red) announced the upcoming stimulus (either heat or sound) and stayed visible throughout stimulation, which was 8 seconds at plateau (roughly 9.5 seconds all in all, depending on calibration). Subjects were then prompted to rate the stimulus. After rating, the white fixation cross reappeared, to turn red again for the next cue. Stimulus modalities were always alternating.

### Sample

A core prerequisite of our analysis strategy was that both modalities (heat and sound) were matched with respect to salience. Although previous studies based estimates of general stimulus characteristics on ratings, such as salience ratings [5] or perceived intensity ratings [13, 15], this can be problematic when comparing modalities, due to differential scaling of responses. We therefore selected skin conductance responses (SCR) as an objective readout parameter linked to modality-independent processes such as emotion, cognition, or attention [24,29–31]. Consequently, our approach is based on comparable SCR for sound and heat stimuli. This necessitated the selection of suitable subjects and experimental sessions that fulfilled this criterion (see Materials and Methods). Analysis included N=26 subjects (50% female, mean age±SD 25.8±3.6; see S1 Tab for more detailed sample characteristics).

### Skin conductance results

As intended by stimulus matching, no significant difference between modalities prevailed (p=0.177) (random intercept model; Fig 2). SCR increased by intensity (t(308)=7.797, p=1e-13). There was no interaction between intensity and modality (p=0.514).

**Fig 2.**
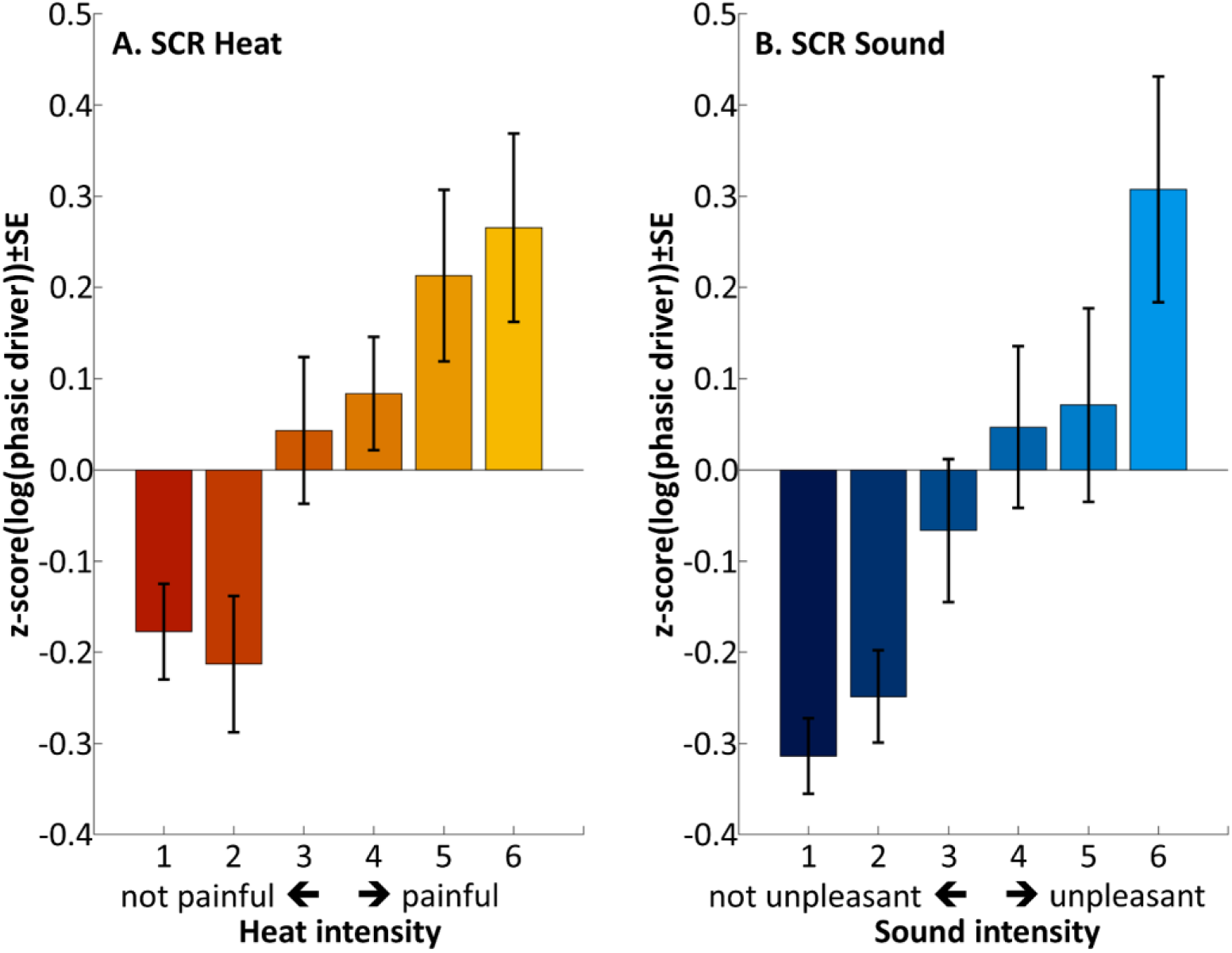
Skin conductance responses following heat (**A**) and sound stimuli (**B**). The pain and unpleasantness thresholds were located between intensities 3 and 4, as per calibration. Individual data is shown in S1 Fig and S2 Fig.

### Stimulus calibration results

Mean heat pain threshold was at 43.3±1.1°C (range 40.5 to 45.4) and corresponded to 50 points on a 0 to 100 point visual analogue scale (VAS). Temperatures for stimulus intensities below and above pain threshold, corresponding to VAS targets of 25, 35, 45, 55, 65 and 75, were 41.7±1.4°C, 42.3±1.2°C, 43.0±1.1°C, 43.7±1.0°C, 44.3±1.0°C and 45.0±1.1°C, respectively. Mean unpleasantness threshold was at 83.9±7.0dBA (range 66.7-99.8). For additional details on heat and sound calibration, see Materials and Methods, and S2 Tab.

### Behavioral results

The analysis of subjective ratings of sound and heat stimuli revealed a significant effect of modality (t(320)=7.820, p=8e-14; average sound rated estimate±SE 13.3±1.7 VAS points higher than average heat) (random intercept model; Fig 3) and a main effect of intensity (t(320)=42.014, p=2e-16; 11.8±0.3 VAS points per intensity step). The interaction between intensity and modality was also significant (t(320)=-4.1529, p=4e-5; 2.3±0.6 VAS points shallower slope in sound, per intensity step).

**Fig 3.**
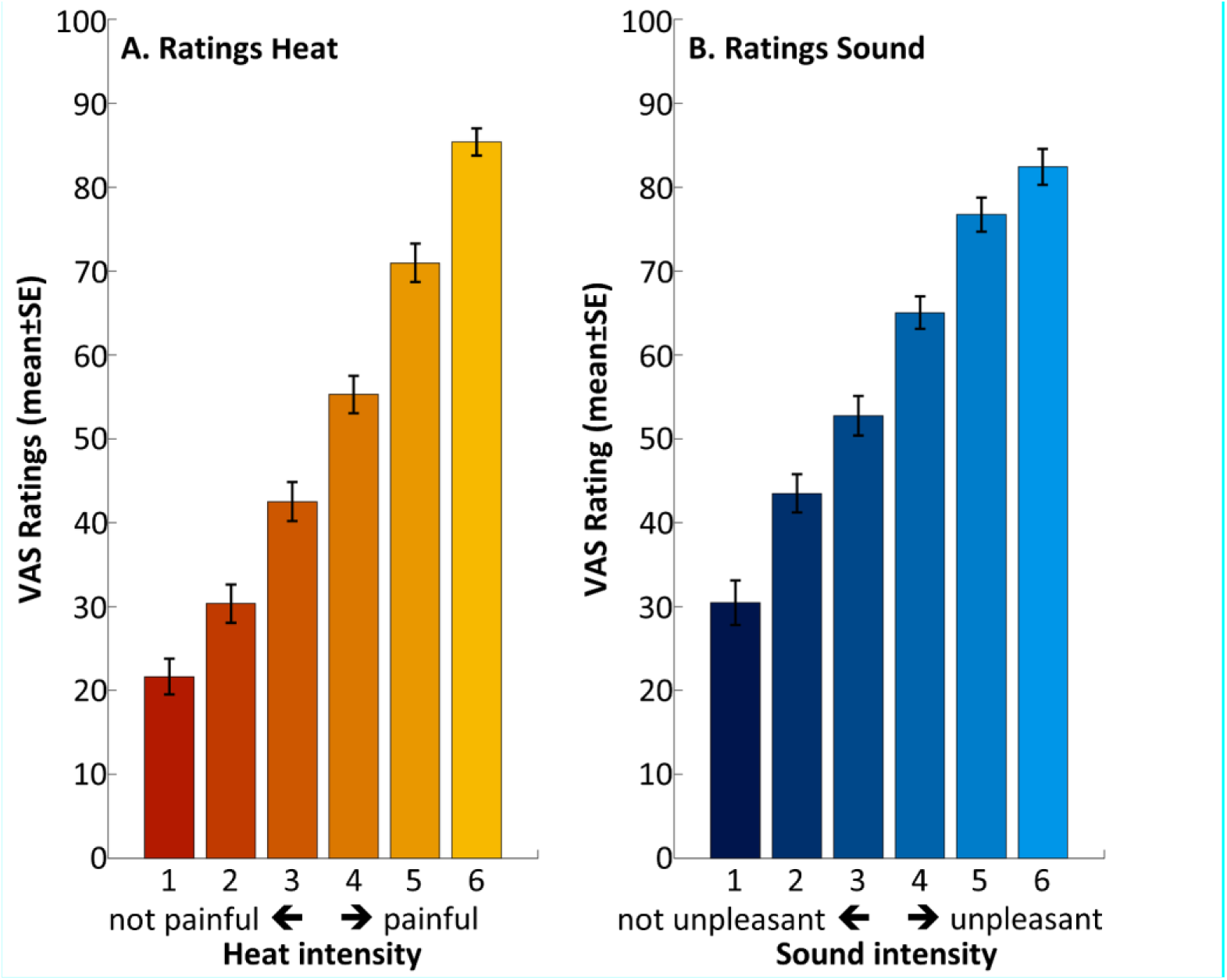
Behavioral ratings following heat (**A**) and sound stimuli (**B**). The pain and unpleasantness thresholds were located between intensities 3 and 4, as per calibration. Individual data is shown in S3 Fig.

### Imaging results

For either modality, a mask was used that was obtained from main effect activations a) larger than the respective comparator modality and b) larger than baseline (S4A Fig; see Materials and Methods for details). The same mask was applied to all contrasts reported in the following, with the exception of conjunction analyses, which were performed without mask. Application of the masks constrains the analyses to areas consistently activated during stimulation of the respective modality. Within the same general area, figures with imaging results use consecutively numbered subscripts (for example, PO_1_ always references the first significant peak described in the parietal operculum).

### Main effects of modality

To test for intermodal differences, we contrasted the main effects for heat and sound (Fig 4). The parietal operculum (secondary somatosensory cortex; peak MNI coordinates x=51, y=-30, z=28, Z=5.62, p(corrected)=1e-05; second peak at x=59, y=-23, z=25, Z=5.221, p(corrected)=1e-04) and dorsal posterior insula (x=40, y=-21, z=19, Z=4.175, p(corrected)=0.012) showed stronger activation for heat as compared to sound. Conversely, Heschl’s gyri (primary auditory cortex; x=64, y=-24, z=7, Z=Inf, p(corrected)=4e-16) showed stronger activation for sound stimuli.

**Fig 4.**
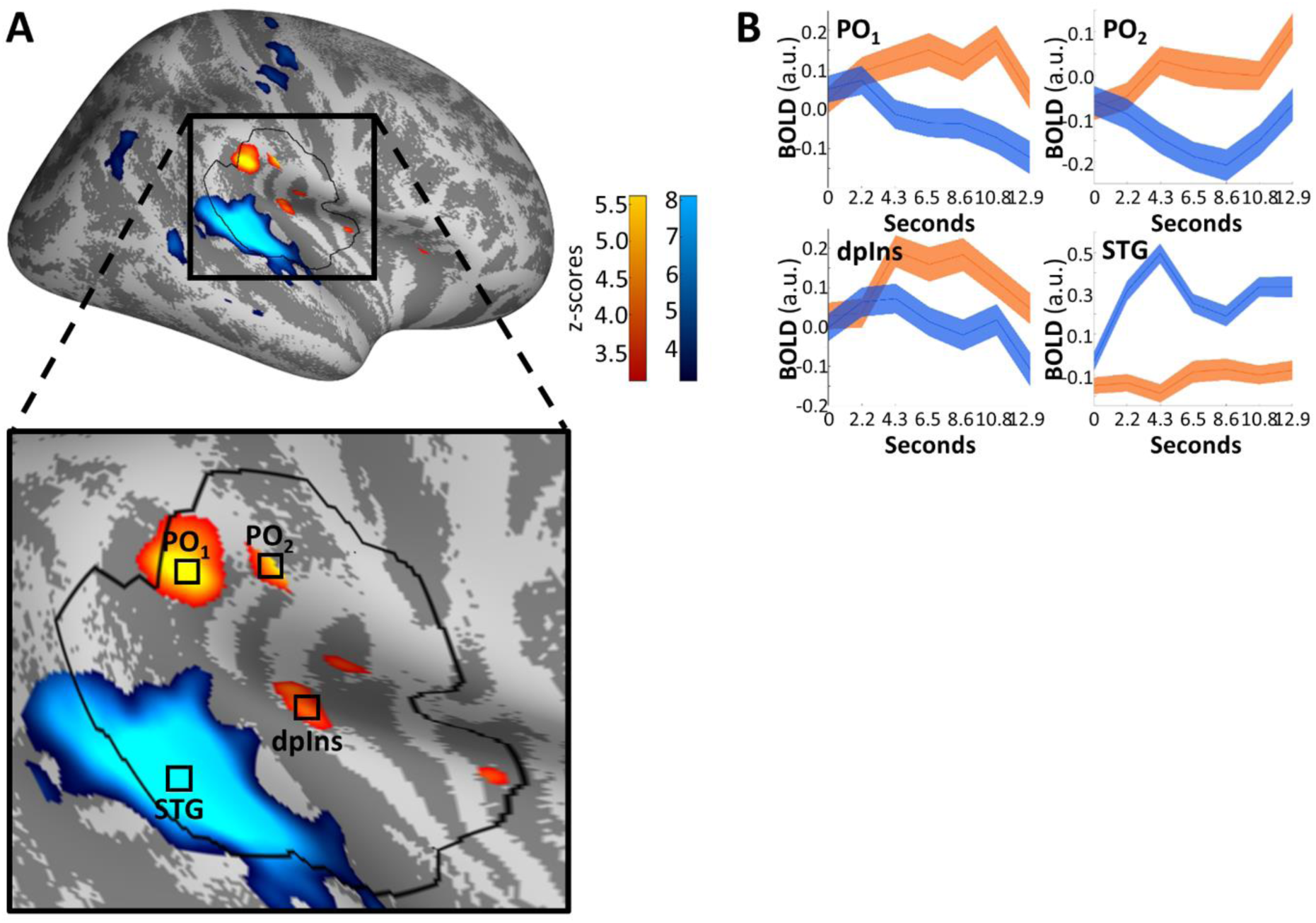
Differential effects of heat (orange) and sound (blue). Significant differences were found in the parietal operculum (PO1 and the more anterior PO2) and dorsal posterior insula (dpIns1) for heat; in the superior temporal gyrus (STG1) and Heschl’s gyri for sound. **A.** Activations are thresholded at p(uncorrected)<0.001 and overlaid on an average brain surface for display purposes. The black line delineates the region of interest used for correction for multiple comparisons. See S5 Fig for peak locations in brain volume slices. **B.** Poststimulus plots of fMRI activation over all stimulus intensities (mean±SE). Subplots PO1, PO2 and dpIns1 show that heat-related activation (orange) dominates in the analyzed time frames (seconds 2.2 through 10.8, see Materials and Methods), while subplot STG1 shows increased sound activation (blue).

Of note, areas activated by either modality show no overlap, as determined via conjunction analyses, even at a liberal threshold of p(uncorrected)<0.001, of contrasts of heat or sound larger than baseline activation. The conjunction analysis did not use any masking; regardless, it did not yield significant results.

### Parametric modulation by stimulus intensity

Irrespective of modality, main effects can be confounded by unspecific effects associated with the generic occurrence of an external stimulus, such as orientation and response preparation. Therefore, we performed an analysis investigating stimulus response functions (SRFs), i.e., testing for stronger BOLD responses for higher stimulus intensities.

We contrasted both modalities to identify areas with diverging SRFs within those areas showing a main effect of either modality, as determined above. For heat, we identified activity in the parietal operculum (x=57, y=-30, z=31, Z=3.999, p(corrected)=0.026) whose SRF diverges from that of the sound modality (Fig 5A). For sound, no significant activity prevailed, that is, no relationship of intensity and brain activity was found within the region of interest. Closer inspection of the time-course of the SRF in the heat modality (Fig 5B) indicates that the SRF’s maximum slope coincides with the peak of the main effect, that is, the modulation of the main effect by intensity is strongest when the main effect itself is strongest.

**Fig 5.**
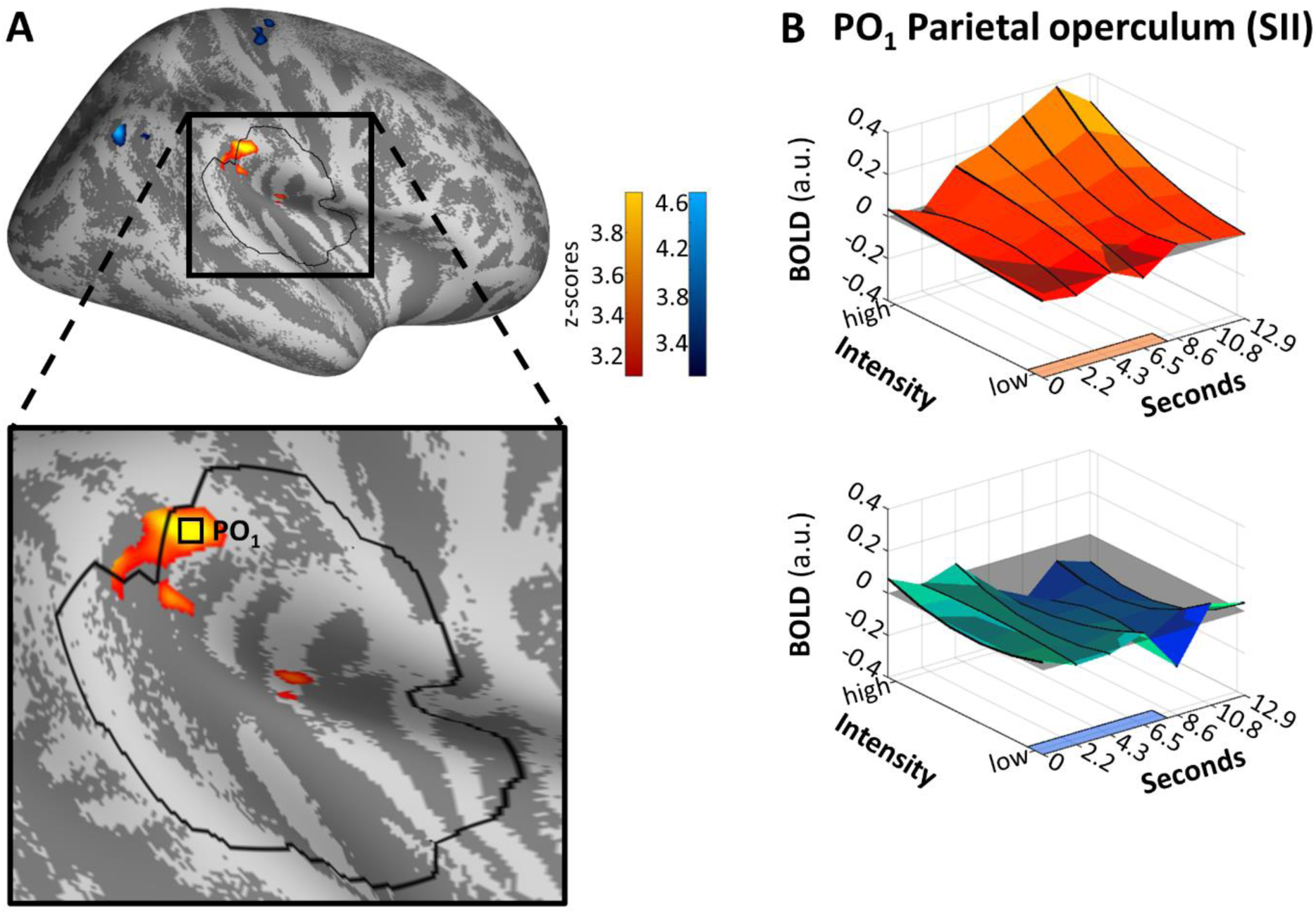
Differential modulation by stimulus intensity for heat (orange) and sound (blue). Significant differences were found in the parietal operculum (PO1) for heat. **A.** Activations are thresholded at p(uncorrected)<0.001 and overlaid on an average brain surface for display purposes. The black line delineates the region of interest used for correction for multiple comparisons. See S6 Fig for peak positions in brain volume slices. **B.** Poststimulus plots of fMRI activation in vertex PO1 during heat (orange) and sound (blue). The colored patches at the right axes show the stimulus duration. The lower left (y-)axes show the parametric modulation affecting the main effect (average size of the effect along the lower right (x-)axes): A straight line parallel to the y-axis indicates no change of the BOLD response depending on stimulus intensity, whereas the a sloped main effect along the y-axis indicates parametric modulation. In this area, the main effect of heat is mostly positively modulated by stimulus intensity, that is, higher stimulus intensities induce a higher extent of BOLD. The highest main effect activation occurs around second 10.8 (corresponding to scan 6), coinciding with the steepest slope of parametric modulation by intensity (y-axis).

Again, an unmasked conjunction analysis at a liberal threshold yielded no significant overlap of both modalities, when comparing contrasts with an SRF slope larger than zero for either modality.

### Parametric modulation by ratings

Although relevant, physical stimulus intensity might not be directly mapped to neuronal activity, as a sensory signal undergoes multiple levels of processing before it reaches cortical areas. We therefore performed an additional analysis, where we investigated whether areas show BOLD responses that are correlated with subjects’ behavioral ratings.

This analysis revealed that activity in the parietal operculum (x=55, y=-37, z=26, Z=5.091, p(corrected)=2e-04; x=58, y=-14, z=18, Z=4.312, p(corrected)=0.008) and the dorsal anterior insula (x=36, y=0, z=14, Z=4.276, p(corrected)=0.009; x=41, y=1, z=14, Z=3.849, p(corrected)=0.047) (Fig 6) showed a positive relationship to perceived intensity. This agrees with and extends results from the previous analysis in which BOLD responses were correlated with stimulus intensity in the parietal operculum. For sound, no significant activity prevailed, that is, no relationship of ratings and brain activity was found.

**Fig 6.**
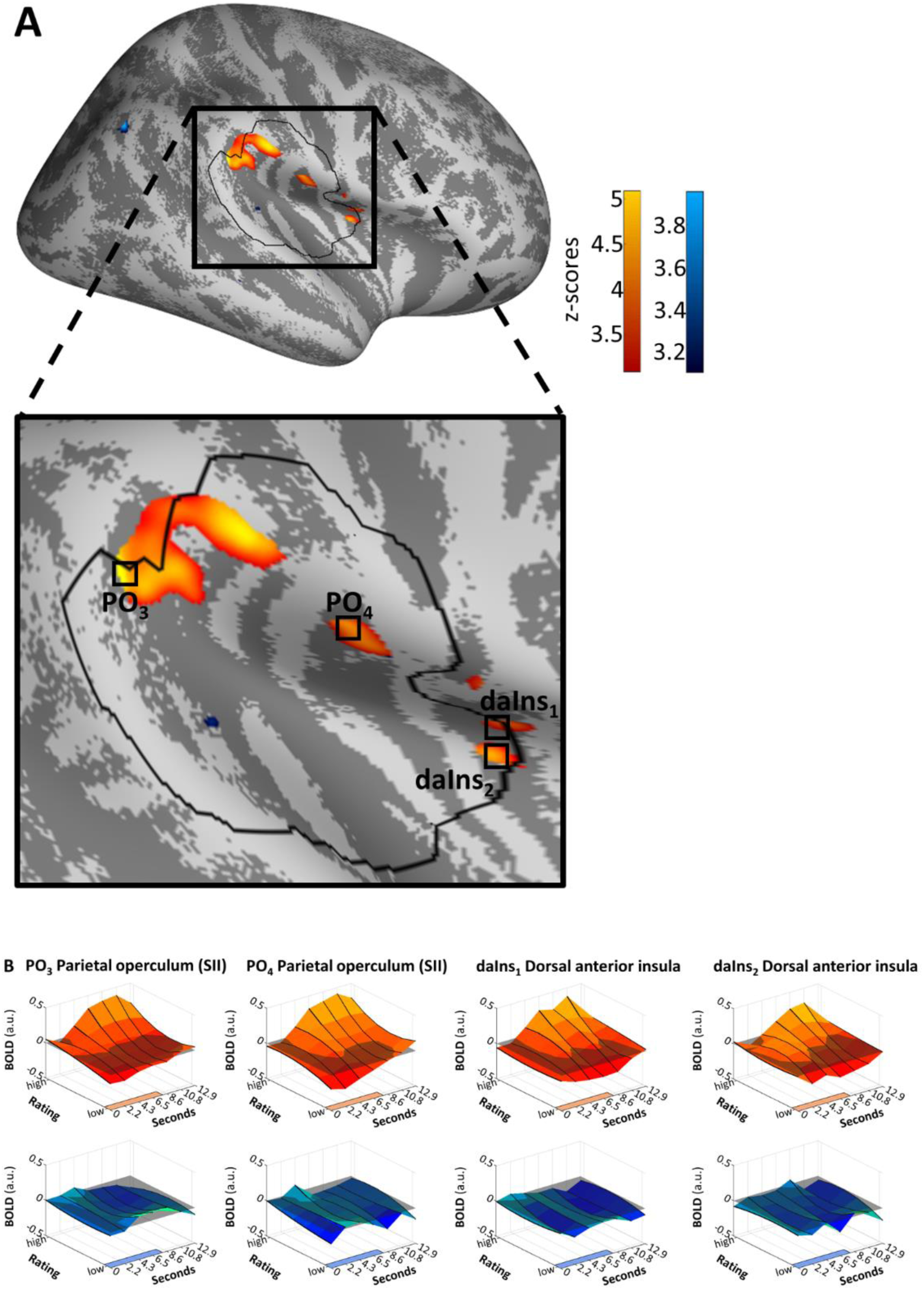
Differential modulation by ratings for heat (orange). Significant differences were found in the parietal operculum (PO3, PO4) and dorsal anterior insula (daIns1 and the more ventral daIns2). **A.** Activations are thresholded at p(uncorrected)<0.001 and overlaid on an average brain surface for display purposes. The black line delineates the region of interest used for correction for multiple comparisons. See S7 Fig for peak positions in brain volume slices. **B.** Poststimulus plots of fMRI activation in vertices PO1, PO3, daIns1 and daIns2 during heat (orange) and sound (blue). The colored patches at the right axes show the stimulus duration. The lower left (y-)axes show the parametric modulation by ratings that are affecting the main effect.

As before, in the unmasked conjunction analysis of contrasts where rating correlated with the BOLD responses, no regions with significant overlap were found.

### Imaging results distinguishing stimuli perceived below and above thresholds

So far, all analyses pooled over non-painful and painful heat percepts. To further investigate pain-related responses, we separated those stimuli reported as non-painful from those reported as painful (i.e., subthreshold versus suprathreshold), and similarly for unpleasant versus non-unpleasant sounds. We followed an axiomatic approach to identify areas where activity under painful stimulation could neither be explained by an overlap with activity under non-painful heat (as would be the case, e.g., in thermosensitive areas), or by an overlap with activity following unpleasant sound (e.g., in areas processing unspecific characteristics of a stimulus). In particular, we posited that a region can be characterized as preferentially pain-processing if the following conditions hold:

▪ Axiom 1: The effect of suprathreshold – i.e., painful – stimulation should be larger than that of subthreshold – i.e., heat – stimulation.
▪ Axiom 2: The effect of suprathreshold heat stimulation should be larger than that of suprathreshold sound stimulation.
▪ Axiom 3: The relationship of ratings and BOLD – i.e., the slope of the stimulus response function – should be stronger for suprathreshold heat than for subthreshold heat.
▪ Axiom 4: The relationship of ratings and BOLD should be stronger for suprathreshold heat than for suprathreshold sound.

Each of the axioms was evaluated at a significance threshold of p=0.05, corrected for family-wise error. After joint application of each axiom, analysis revealed activation in the posterior parietal operculum (x=56, y=-37, z=25, Z=3.909, p(corrected)=0.035) (Fig 7), adjacent to the supramarginal gyrus. To broaden the scope, we have performed the same axiomatic analysis on the whole surface with a lower significance threshold of p(uncorrected)=0.001 (S8 Fig). For comparison, results from a conventional 3D analysis are also provided for the whole brain, again with a lower threshold of p(uncorrected)<0.001 (S9 Fig).

**Fig 7.**
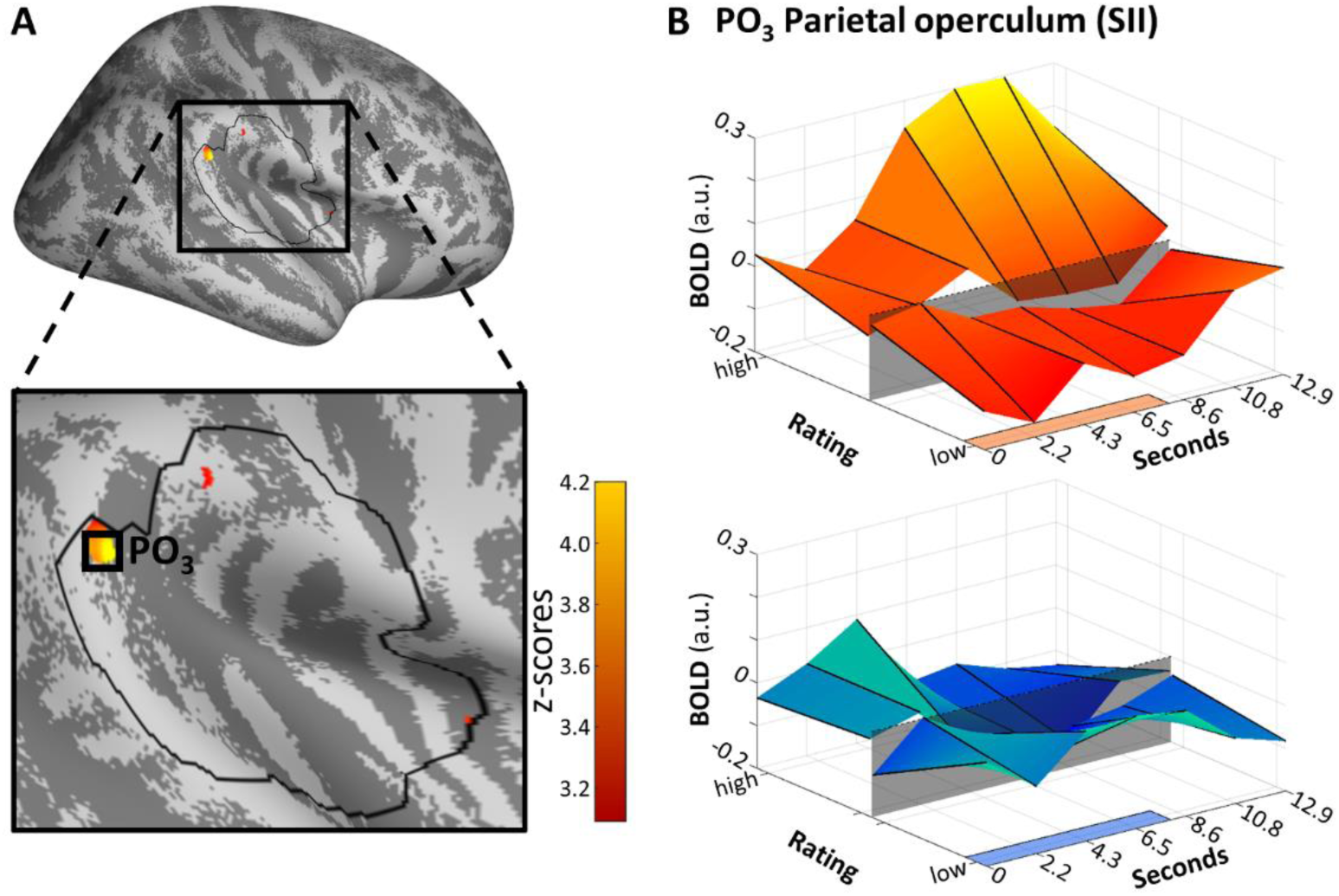
Areas that fulfill the axiomatic requirements of differential activation during pain compared to heat and sound. In detail, these axioms were 1) a larger effect of suprathreshold heat compared to subthreshold heat, 2) a larger effect of suprathreshold heat compared to suprathreshold sound, 3) a stronger relationship of BOLD with pain ratings than with heat ratings, 4) a stronger relationship of BOLD with pain ratings than with unpleasantness ratings. Significant activation was found in the parietal operculum (PO3). **A.** Activations are thresholded at p(uncorrected)<0.001 and overlaid on an average brain surface for display purposes. The black line delineates the SVC mask. See S10 Fig for peak positions in brain volume slices. **B.** Poststimulus plots of fMRI activation in vertex PO3 during heat (orange) and sound (blue). The shaded patch in the center signifies the pain threshold (for heat) and unpleasantness threshold (for sound). The colored patches at the right axes show the stimulus duration.

## Discussion

This study aimed to identify regions relevant for heat pain processing, and to determine whether their activation can be explained by salience or other modality-independent characteristics. We used individually calibrated, parametrically graded heat stimuli, and an auditory control condition. Heat and sound stimuli were matched for arousal as indicated by similar skin conductance responses. Furthermore, we employed surface-based analyses to mitigate spatial inaccuracies of 3D smoothing.

Main effects for heat were identified in the parietal operculum and the posterior insula, main effects of acoustic stimuli were observed in the superior temporal gyrus. More importantly, in the parietal operculum, we observed a differential correlation of brain activity with ratings above versus below the heat pain threshold, concurrent with a differential correlation with ratings under painful heat versus unpleasant sound. As we have matched both modalities for salience, these results rule out, within the limitations of this experimental approach, that activity in this area is simply related to stimulus salience, and suggests a more dedicated role in heat pain processing.

Using SCR as an autonomic readout of arousal [24,29–31] allowed us to establish comparable salience of the stimulus material, independent of any behavioral assessments. Although salience can be assessed psychometrically [5] and research exists to establish concurrent validity of salience ratings within individual modalities [32], to our knowledge, such ratings have not been validated cross-modally. It is likely that salience ratings are scaled differently according to some modality-specific perceptual range, that is, a rating of X on a “salience” scale while assessing pain may mean something different from a rating of X while assessing sound, identical scale anchors notwithstanding. To our knowledge, a “ground truth” to compare multiple sensory modalities has never been established in this regard. Our results support the notion of modality-specific scaling, as we have observed a prevailing difference in behavioral ratings between the two modalities (sound was, on average, rated as more aversive, but had a shallower slope with increasing intensities). This means that a reliance on behavioral ratings alone could compound SCR dissimilarities between comparator modalities.

With six graded stimulus intensities per modality, our design allowed for the assessment of stimulus response functions as opposed to simple mean comparisons between a single intensity and a low-level baseline, or between single sub- and suprathreshold stimuli. Apart from physical intensities, this also allowed us to use a large range of individual ratings as predictors. Using these perceived intensities, we were able to directly investigate competing modes of encoding. For example, a brain area may encode heat intensity, regardless of nociceptive intensity, or it may be inactive below threshold but encode pain intensity above threshold [8,20,21]. In the analysis distinguishing between sub- and suprathreshold stimuli, we see a clear pain intensity-related response in the parietal operculum (Fig 7). While not a main focus of this paper, we do see a shift in SRFs even within small cortical distances: For example, an area rostral (x=55, y=-26, z=26) to the posterior parietal operculum cluster identified in the axiomatic analyses (x=56, y=-37, z=25) fails to register differences in the parametric modulation by sub-versus suprathreshold heat (Fig 8).

**Fig 8.**
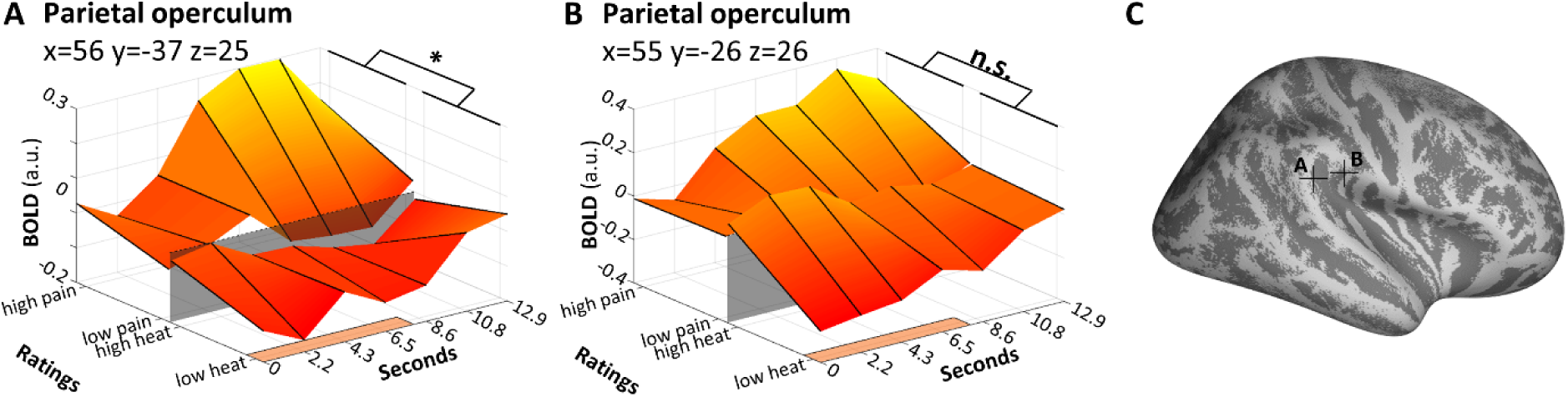
Distinction of areas with stimulus response functions corresponding to the axioms, or not (heat modality only). **A.** The slopes of subthreshold (heat) versus suprathreshold (pain) activation as described in Fig 7, corresponding to parietal operculum cluster 3 (PO3). Heat slopes are shallower than pain slopes. **B.** Slopes of heat and pain activation in an opercular vertex slightly anterior to A (corresponding to PO2), as determined per conjunction of heat and pain parametric modulation. Slopes are more aligned, preventing the contrasted activation (pain>heat) of reaching significance. Note that this is not a formal comparison to A. **C.** Location of the vertices described in A and B. B is in an adjacent area about 1 cm surface distance rostral from A.

Areas in the insula and surrounding cortical areas are characterized by extreme cortical folding. We therefore implemented a subject-specific surface-based fMRI analysis. This prevents contamination of gray matter voxels by signal from white matter and cerebrospinal fluid. Furthermore, surface-based analyses circumvent potential issues arising from three-dimensional smoothing which accidentally mixes signals from structures adjacent in three-dimensional space which are actually distant from each other. For example, the parietal operculum is directly adjacent to the superior temporal gyrus in three-dimensional space, but their neurons are separated by the entire insular fold. Smoothing with a three-dimensional kernel therefore includes activity across the lateral sulcus (alongside noise from white matter and corticospinal fluid), thereby increasing error terms and decreasing sensitivity of the respective comparisons. Importantly, in this case, three-dimensional smoothing could also generate erroneous overlaps between conditions. Surface-based analyses have been found to increase sensitivity and reduce deviations when normalizing from native to standard space [33].

In contrast to previous multimodal studies, we explicitly chose modalities where the aversiveness would be generated by virtue of physical intensity. This is naturally the case with painful stimulation, but several studies did not use aversive stimulation in non-painful control modalities (for example [11], who used low-intensity tactile stimulation). Consequently, the acoustic modality was chosen because stimuli can be generated in close analogy to heat, by altering the physical intensity of the stimuli.

For our analyses of neuronal activity, we have focused on the posterior insula and adjacent areas. The insula is of particular interest, because its involvement in pain has been well-documented [4,11,34]. Furthermore, it has been hypothesized to perform polymodal magnitude estimation [8, 35], and is also involved in threat learning [36] and salience processes [37, 38]. Unambiguous data concerning the involvement of the insular cortex in pain processing also comes from direct cortical stimulation studies [39–41]. Consequently, it is a prime candidate to assess overlaps and differences in activation patterns.

With its reliable activation following painful stimulation and within the constraints inherent in our experimental approach, we can corroborate the role of the parietal operculum as an important area of heat pain processing, and importantly, one whose activity cannot be explained by unspecific characteristics of sensory stimulation, like stimulus salience. The area not only shows increased activation when comparing pain and other modalities (heat, sound), but also exhibits a monotonic increase with perceived pain. The peak of the BOLD response following pain clearly coincides with the largest modulation by behavioral ratings of pain, roughly 8 seconds after stimulus onset (Fig 7B). Importantly, this area has close functional connections with the posterior insula [42], another area of interest [11, 25]. Contextualization of this finding with previous studies comparing at least two stimulus modalities [5,8,13,43] is difficult, because their modality-matching was performed on ratings of salience and/or intensity (not SCR), or used only a single stimulus intensity per modality (i.e., a comparison of main effects). With these caveats in mind, one study found activation at similar coordinates for painful stimulation (x=56, y=-36, z=28) and the visual control condition (x=62, y=-36, z=20), but no preferential activation following pain [8]. Mouraux and colleagues [5] likewise reported no preferential S2 activation following painful stimulation, but an increased activation during (noxious and non-noxious) somatosensory stimulation compared to acoustic and visual. Another study reanalyzed two studies (including [5]) and identifies a proximate cluster (x≈53, y≈-27, z=27) as “pain-preferring” in the sense that activation was consistently higher compared to all three control modalities [43]. The last study used a tactile control condition and did not report preferential activation of the cluster following painful stimulation [13].

Previous studies have shown a considerable overlap of brain activity following stimulation in different sensory modalities [5,13,14]. The extent of any overlap in functional neuroimaging is strongly dependent on the statistical threshold employed, and it is possible that diverging results stem from different thresholding or conjunction methods. However, for reasons of comparability we wish to mention that our data shows no such overlap of activation across modalities, even at a low threshold. While other differences in imaging analysis exist between the studies (for example, we have performed analysis on the brain surface), the lack of overlap might rather be related to the differences in stimulus parameters: Previous multisensory studies have used rapid onset stimuli of very short duration [5,13,14], whereas ours were considerably longer (8 seconds plateau, circa 9.5 seconds with upward/downward slopes). It is possible that with increasing brevity and suddenness of the stimuli, the extent of unspecific orientation responses and other attention related processes is disproportionally larger, and therefore a larger overlap of neuronal activation can be observed [44]. If true, this overlap would naturally be determined, to a large extent, by unspecific and not pain-related activations such as salience.

Brain responses evoked by stimuli in different sensory modalities might follow different time courses; systematic differences may, for example, arise from different conduction speeds of fibers relaying auditory (mostly very fast Aα fibers), thermoceptive (mostly slow C fibers) and nociceptive input (Aδ and C fibers), compounded by the fact that thermal stimulation (in this setup) occurs at a distal site compared to auditory stimulation. Therefore, in analogy to [5], we opted for analyzing the time-course of all imaging data by using finite impulse responses as basis functions. This largely avoids the constraints and biases implicit in comparing mean activations obtained by pre-defined hemodynamic response functions.

Some limitations apply to the present research. Importantly, due to the salience-matching approach based on skin conductance, a substantial number of study participants were not eligible for inclusion, because they either did not show a stimulus-intensity related SCR, or showed responses skewed towards the heat modality. This procedure could induce a sampling bias, for example by selecting subjects with stronger SCR reactivity, which could implicate different salience network connectivity [45]. However, we found no difference in calibrated intensities in pain or sound or their variance when comparing included versus excluded subjects (all p>0.5; see S3 Tab), making a sampling bias unlikely.

As additional limitation, the study used stimuli of mild to moderate aversiveness (calibrated to a maximum of 50 if rescaled to a conventional, 0-100 suprathreshold visual analogue scale). This aspect, too, could be amended to cover a broader range, albeit increasing the risk of carry-over effects such as sensitization, particularly with longer stimulus duration. Furthermore, the use of only a single trial-based, post-stimulus rating of stimulus intensity could be criticized. In fact, one common recommendation for pain measurement is to distinguish multiple pain dimensions [1], most frequently intensity and unpleasantness [46], although these aspects tend to be highly correlated in non-interventional designs [47, 48]. Given the SCR-based approach to equalize salience and to include more stimulus repetitions, we opted against multiple VAS for protocol reasons, namely ease of measurement and to avoid confusion.

While we have identified areas preferentially active in painful heat as compared to unpleasant sound, we cannot claim that these areas are specific for pain. In fact, it is important to note that specificity cannot be ascertained with a limited number of control conditions [19, 49]. We concur that the notion of specificity is more academic in nature than might benefit the field [3]. The preferences of certain areas to process various inputs – whether visual, acoustic, nociceptive – is best construed as a matter of degree, that is, a question of specialization rather than specificity, as has been suggested for functions unrelated to pain [50]. Nevertheless, the rigorous axiomatic approach allows for a strong hypothesis ascribing the parietal operculum a dedicated role in pain processing.

## Materials and Methods

The protocol was approved by the local Ethics Committee (Ethikkommission der Ärztekammer Hamburg, vote PV4745) and conformed to the standards laid out by the World Medical Association in the Declaration of Helsinki. Participants gave written informed consent prior to participation.

### Exclusion criteria

A list of exclusion criteria is provided in Table 1.

**Table 1.**
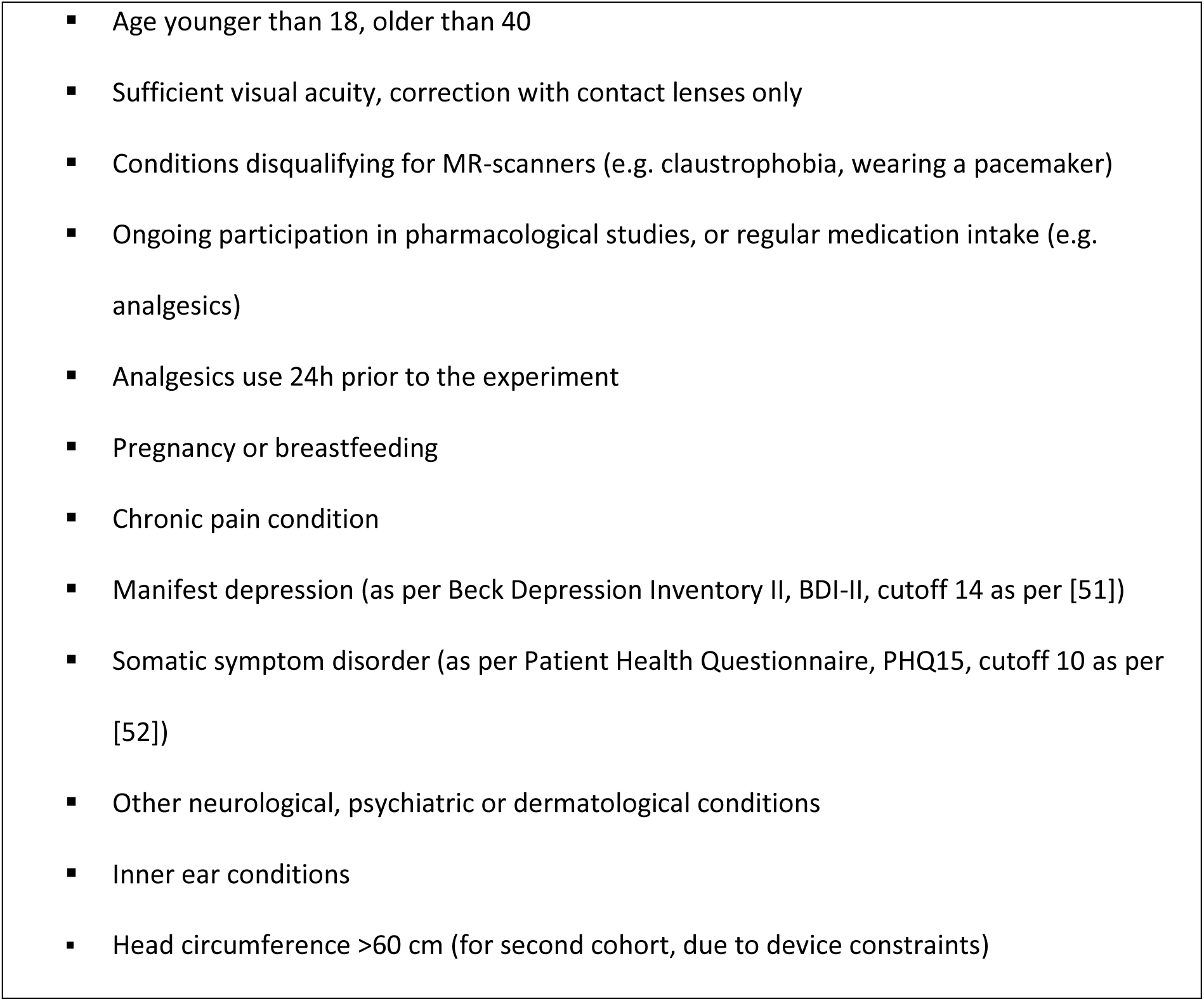
Exclusion criteria.

### Psychophysiological recordings

Electrodermal activity was measured with MRI-compatible electrodes on the thenar and hypothenar of the left hand. Electrodes were connected to Lead108 carbon leads (BIOPAC Systems, Goleta, CA, USA). The signal was amplified with an MP150 analog amplifier (also BIOPAC Systems). It was sampled at 1000 Hz using a CED 1401 analog-digital converter (Cambridge Electronic Design, Cambridge, UK) and downsampled to 100 Hz for analysis.

Analysis was performed using the Ledalab toolbox for MATLAB [53]. Single subject data were screened for artifacts which were removed if possible by using built-in artifact correction algorithms. Using a deconvolution procedure, we computed phasic skin conductance (SCR). SCR occurring after stimulus onset and within stimulus duration was used for measuring autonomic arousal. Response windows were defined by visual inspection, per modality: between 2.0 s and 4.5 s for heat, and between 1.5 s and 4.0 s for sound. Results were log- and z-transformed to reduce the impact of intra- and interindividual outliers [24]. Subsequently, SCR was averaged within subjects for two modalities (heat/sound) and six stimulus intensities each, yielding twelve values per person.

SCR was used because it is an objective measure of general sympathetic activity, and therefore a measure of arousal, stimulus salience and several associated psychological processes [24,29–31]. It is is routinely used in assessing painful [23,54,55] as well as acoustic stimulation [56].

### fMRI acquisition and preprocessing

Functional and anatomical imaging was performed using a TRIO 3T MR Scanner (Siemens, Erlangen, Germany) with a 12-channel head coil. An fMRI sequence of 36 transversal slices of 2 mm thickness was acquired using T2*-weighted gradient echo-planar imaging (EPI; 2150 ms TR, 25 ms TE, 80° flip angle, 2×2×2 mm voxel size, 1 mm gap, 216×216×107 mm field of view, acceleration factor of 2 with generalized autocalibrating partially parallel acquisitions reconstruction, GRAPPA). Coverage did not include the apical parts of the frontal/parietal lobes. Additionally, a T1-weighted MPRAGE anatomical image was obtained for the entire head (voxel size 1×1×1 mm, 240 slices).

For each subject, fMRI volumes were realigned to the mean image in a two-pass procedure, and co-registered to the anatomical image using affine transformations. Anatomical images were segmented into tissue types, and individual brain surfaces generated, using the CAT12 toolbox for SPM (Christian Gaser & Robert Dahnke, http://www.neuro.uni-jena.de/cat/).

### Analysis of imaging data

Subject-level analyses were performed on the 3D (volume) data in native space without smoothing, using an implicit mask at 0.6 to facilitate subsequent (surface) processing. We computed general linear models to identify brain structures involved in the processing of each stimulus modality, as well as the encoding of intensities within those modalities. All analyses were performed with seventh order FIR basis functions, of which bins 2 to 6 are considered when comparing conditions. This amounts to seconds 2.2 through 10.8 post stimulus onset. Realignment (motion) parameters as well as regressors obtained from ventricular motion were included as nuisance variables, to mitigate motion-related artifacts.

We first set up a model including one regressor for stimulus main effects in each modality. Another two regressors – one linear, one quadratic – encoding stimulus intensities 1 through 6 were added per modality, as parametric modulators. The second model likewise included main effects, and behavioral ratings as linear and quadratic parametric modulators. Finally, the third model further distinguished the two modalities in stimuli perceived as below and above the respective thresholds (pain for heat stimuli, unpleasantness for sound stimuli), yielding four main effect regressors (subthreshold heat, suprathreshold heat – i.e., pain –, subthreshold sound, suprathreshold sound). Behavioral ratings were again included as linear parametric modulators; quadratic modulation was not considered to preclude overfitting.

Results from subject-level analyses were mapped to brain surfaces obtained via the CAT12 segmentation procedure. The mapped subject-level results were then resampled to correspond to surface cortical templates, and smoothed with a 6 mm full width-half maximum 2D kernel. Group-level analyses were performed including the mapped contrasts, which are described in the Results section.

Masking was used to distinguish either modality, as the ANOVAs employed are unsigned and in principle detect differences in activation regardless of direction. Therefore, we obtained signed (that is, unconstrained by p values) masks from calculating a conjunction from significant voxels of a) a t-test contrasting the average main effects of either modality (i.e., where activation following heat was larger than that following sound, and vice versa), and b) a t-test contrasting either modality to low-level baseline (i.e., where activation following heat – or sound, respectively – was larger than zero. This yielded a single mask for both modalities, which was applied to all analyses (unless otherwise noted) (S1B Fig).

For the purpose of this study, we focused on the hemisphere contralateral to the stimulation, in our case the right hemisphere. In general, the larger part of activity following pain is contralateral to the stimulation site, but is known to be bilateral in several key areas such as the secondary somatosensory cortex and the insula [22].

Furthermore, we focused on the insula and directly adjacent areas for small volume correction of significance level. In particular, we included the granular insular cortex (Ig1, Ig2) as well as the parietal operculum (OP1, OP2) and primary auditory cortex (Te1.1), using the SPM Anatomy Toolbox (version 2.2b [57]). This mask was mapped to a template brain surface, then smoothed with a 4 mm 2D kernel to close gaps. The resulting binary mask (S1A Fig) was roughly centered around previously reported coordinates (x=[-]34, y=-20, z=18) involving areas putatively dedicated to pain processing [11]. It was used for small volume correction of second level analyses, where results were considered after correction for family-wise error rate of p<0.05.

### Psychometry

Owing to the study’s aim to compare two stimulus modalities (heat and sound), they had to be presented and rated in an analogous fashion. Therefore, while retaining the intuitive descriptor “painfulness” for rating noxious heat (as composite measure of intensity and unpleasantness), we settled on “unpleasantness” as descriptor for sounds. This also seemed warranted considering the high correlation of intensity and unpleasantness measures in heat pain [48], while unpleasantness is one of the definitional criteria of pain [2].

Furthermore, since we wanted to use graded stimuli both below and above the respective thresholds (pain threshold for heat, unpleasantness threshold for sound), we deviated from the more common simple visual analogue scales (VAS) and devised two partitioned 0 to 100 VAS for both modalities (Fig 1A).

For heat, it captured both painful and non-painful sensations. Subjects were instructed to indicate heat intensity in absence of pain in the 0 through 49 range, and heat pain intensity in the 50 through 100 range. Hence, anchors were displayed for “no sensation” (0), “minimal pain” (50), and “unbearable pain” (100). Pain was operationalized as the presence of sensations other than pure heat intensity, such as stinging or burning, as per the guidelines of the German Research Network on Neuropathic Pain [58].

Likewise, for sound, both unpleasant and non-unpleasant sensations were captured by the VAS. Subjects were instructed to indicate loudness in absence of unpleasantness in the 0 through 49 range, and loudness unpleasantness in the 50 through 100 range. Anchors were displayed for “inaudible” (0), “minimally unpleasant” (50), and “extremely unpleasant” (100). Unpleasantness was operationalized as a bothersome quality of the sound emerging at a certain loudness.

### Heat stimuli and calibration

Heat stimuli were delivered using a CHEPS thermode (Medoc, Ramat-Yishai, Israel). Stimulation sites were located on the radial surface of the forearm. Three separate sites were used for calibration and either experimental session, to avoid changes in heat/pain perception due to repeated stimulation. Around the middle of the forearm (half distance between crook of the arm and distal wrist crease; see Fig 1B), three stimulation sites were marked prior to the experimental sessions. For calibration, a medial site on the distal part of the forearm was used; for experimental sessions 1 and 2, two adjacent proximal sites were used, in counterbalanced order. During both calibration procedure and experimental sessions, baseline temperature was set to 35°C, and rise and fall rate were set to 15°C per second. The duration of heat stimuli was set to eight seconds at target temperature (plateau), except for preexposure stimuli whose plateau duration was zero (and thus only consisted of temperature up- and downramping).

A two-step stimulus calibration was performed for each subject, to determine three temperatures below the individual pain threshold, and three above. Calibration was performed with the MR-scanner running the same sequence as during the actual experimental sessions, to mimic ambient conditions [59]. fMRI data from calibration was later discarded.

In a first calibration step, the pain threshold was determined. Subjects were preexposed to four brief heat stimuli. Preexposure started at 42°C and each consecutive stimulus was increased by 0.5°C, up to 43.5°C. If a subject indicated the last stimulus as painful, starting temperature for the following procedure was set to 43°C, else to 44°C. We then used a probabilistic tracking procedure for threshold determination, assuming a normal distribution of pain perception around the actual threshold [60]. Eight full-length stimuli were presented and received a binary rating (painful or not painful). Depending on the rating of the previous stimulus, each consecutive stimulus was set to a higher or lower temperature according to the probability informed by previous pivot points. The final temperature was defined as threshold intensity.

In a second calibration step, eight stimuli unevenly spaced around threshold intensity (from -2°C to +1.6°C, with smaller intervals towards ±0°C) were rated on the partitioned VAS described above. After the procedure, linear regression was used to calculate target temperatures H1 through H6, to obtain subthreshold VAS ratings of 25, 35 and 45 (H1–H3), and suprathreshold VAS ratings of 55, 65 and 75 (H4–H6).

These six intensities were used throughout the experimental sessions.

### Sound stimuli and calibration

Sound stimuli were delivered using MR-compatible headphones (NordicNeuroLabs, Bergen, Norway). A pure sound (frequency 1000 Hz, sampling rate 22050 Hz) was generated using MATLAB. A log function was used to translate increases in (physical) amplitude to smooth gradual increases in (psychoacoustic) loudness, to mimic the heat stimuli’s temperature ramps. Like the heat stimuli, sound stimuli were presented for eight seconds at target loudness (plateau), and the scanner was running a dummy EPI sequence throughout to mimic actual conditions [61].

A two-step stimulus calibration was performed for each subject, to determine three sounds below the individual unpleasantness threshold, and three above. The general procedure was analogous to the one used for heat.

In a first calibration step, the individual loudness unpleasantness threshold (in percent of maximum amplitude of ∼100 dB, allowing for safe exposure even at maximum intensities [62]) was determined by an ascending methods of limits-procedure. Six sounds of gradually increasing loudness were played. The calibration sounds differed in the steepness of the loudness ramps, taking between 9 and 15 seconds to reach peak amplitude. Subjects were asked to indicate the point where the loudness became unpleasant. The mean of the last four of the six stimuli was defined as threshold loudness.

In a second calibration step, 16 stimuli unevenly spaced around threshold loudness were presented (from -15% to +15%, smaller intervals towards ±0%), with stimulus characteristics set to mimic those of heat stimuli (roughly 0.75 s ramps up and down, plus 8 s plateau loudness). As with heat ratings, linear regression was used to calculate target amplitudes S1 through S6, namely to obtain subthreshold VAS ratings of 25, 35 and 45 (S1–S3), and suprathreshold VAS ratings of 55, 65 and 75 (S4–S6). For the second cohort, VAS targets were informed by the corresponding mean SCR amplitude of the first cohort (see “Differences between first and second cohort”).

Finally, ramping characteristics of sound stimuli (the seconds it took to plateau) were set to correspond to those of the respective intensity’s heat stimuli, such that corresponding intensities of both modalities had an identical overall length (ramps plus plateau).

### Stimulus presentation during experimental sessions

After calibration, the thermode stimulation site was changed, and the first experimental session commenced. Heat and sound stimuli were presented in alternation, so that trials of the same modality were spaced with an intertrial interval of approximately 30 seconds. Each trial followed the same basic structure (Fig 1C).

Within each modality, the six intensities were pseudorandomized in microblocks. Randomization was performed such that each sequence of six stimuli contained one instance of each intensity. It was further constrained such that the very first stimulus was never chosen from the highest two intensities, and two consecutive intensities were never more than 3 intensity steps different (e.g., the intensity following heat intensity 1 could not exceed heat intensity 4).

After changing thermode stimulation site again, session 2 commenced with identical protocol (albeit different randomization).

Visual cues and VAS rating scales were displayed in the scanner using back-projection via a 45° mirror placed atop the head coil.

### Selection of subsample for analysis with comparable SCR between modalities

In total we assessed two cohorts of 32 subjects and 26 subjects. To obtain an “SCR-equalized” subsample from all subjects (N=58 with 2 sessions, that is a total of 116 experimental sessions), in a first step, we excluded all sessions where the correlation between ratings (that is, perceived stimulus intensity/unpleasantness) and SCR was lower than or equal to zero, so that only subjects with a positive correlation in both modalities were eligible for the next step.

In a second step, we used Bayes factors [63, 64] to determine the flipping point where modality became obsolete as explanatory variable. Bayes factors express the ratio of the marginal likelihood of the data under the compared models; since they consider the number of free parameters, they allow for the selection of the “better” model (best fit to the data and most parsimonious). For every session, we obtained the mean SCR (log-transformed and normalized values) for both modalities; sessions with the largest predominance of heat-SCR were then consecutively removed. After each removal, we obtained the Bayes factors for the remaining sample, comparing the model with intensity only as predictor to that with modality added as predictor. Once the Bayes factor dropped below 1 (meaning that the addition of modality as predictor did not serve to improve the model), we stopped the pruning procedure. This relatively permissive criterion for session inclusion was chosen in order to preserve as many sessions as possible.

This procedure yielded a sample where modality did not contribute to explaining the SCR data, with 26 unique subjects contributing 33 sessions. From the first cohort, 15 subjects contributed 19 sessions, from the second cohort, 11 subjects contributed 13 sessions to the SCR-equalized analysis.

### Differences between first and second cohort

Since we had determined that not every person’s skin conductance responded to both modalities to a comparable extent, we set out to select a subsample of persons who had comparable SCR. To reach a sufficient number of such “responders”, we had to perform an additional data collection.

Because of logistical reasons (scanner upgrade in January 2018), some parameters of fMRI acquisition had to be modified for the new PRISMA 3T MR Scanner (Siemens, Erlangen, Germany). Instead of a 12-channel head coil, we had to employ a 20-channel head coil. Delivery of the auditory stimulus was performed with a CONFON headphone (Cambridge Research Systems Ltd, Rochester, United Kingdom). These measures necessitated the exclusion of subjects with head circumference above 60cm.

Furthermore, to facilitate increased SCR responding to sound, we increased the amplitude of the sound stimuli. Using calibration data from the first data collection and linear extrapolation, we calculated sound VAS targets required to induce SCRs of an amplitude comparable to those of heat VAS targets of the same intended intensity 1 through 6. We determined that corresponding to our heat VAS targets of 25, 35, 45, 55, 65, 75 (see “Heat stimuli and calibration”), we would need to apply sound amplitudes inducing sound VAS targets of 48, 59, 70, 82, 93, 105. Furthermore, during subject instruction, we emphasized the fact that the amplitude of sound stimuli was not within pathological range. This was done to prevent overly cautious subject behavior, following anecdotal evidence from the first cohort that sound stimuli were associated with higher safety concerns than heat stimuli.

### Statistical analyses

All analyses were performed using MATLAB (version R2017b) and SPM12 (version 6906). Significance level was set to p=0.05 for psychophysiological and behavioral data, whereas imaging results were corrected using family-wise error rate adjustment at p<0.05. For visualization, activations are thresholded at p(uncorrected)<0.001 and overlaid on an average brain surface. All coordinates are reported in Montreal Neurological Institute (MNI) space.

Skin conductance data and behavioral ratings were analyzed using linear mixed models with random intercepts [65], with centering of predictors following recommendations [66].

Group-level analyses of imaging data were performed as within-subjects ANOVA (cf. “Analysis of imaging data” above, and the respective Results sections).

## Acknowledgements

This work was supported by European Research Council Advanced Grant ERC-2010-AdG_20100407 and Deutsche Forschungsgemeinschaft Grant SFB 936 Project A06. Björn Horing was supported by the Alexander von Humboldt-Foundation (Feodor Lynen Return Fellowship). We thank Jürgen Finsterbusch, Katrin Bergholz, Waldemar Schwarz and Kathrin Wendt for technical assistance during MR data collection, and Lara Austermann, Tim Dretzler, Katharina Ebel and Matthias Kerkemeyer for their assistance with data collection, and finally Christian Gaser for helpful comments concerning the CAT12 toolbox.

## Supporting information

**S1 Tab.** Sample descriptive statistics.

| Questionnaire | Construct | Mean±SD | Sample range | Possible range |
| --- | --- | --- | --- | --- |
| **BDI-II** [SR 1,2] | Depression | 2.0±2.2 | 0-8 | 0-63 |
| **PHQ15** [SR 3] | Somatization | 3.9±2.3 | 0-9 | 0-30 |
| **FPQ** [SR 4] |  |  |  |  |
| severe | Fear of pain | 28.9±8.3 | 11-42 | 10-50 |
| minor | Fear of pain | 14.0±4.5 | 10-27 | 10-50 |
| **PVAQ** [SR 5] | Pain vigilance and awareness | 34.6±8.7 | 21-54 | 0-80 |
| **PSQ** [SR 6] | Pain sensitivity | 45.1±14.8 | 17-73 | 0-140 |
| **PRSS** [SR 7] |  |  |  |  |
| Catastrophizing | Pain catastrophizing | 8.6±5.4 | 2-21 | 0-45, higher more catastrophizing |
| Coping | Pain coping | 29.8±5.2 | 19-38 | 0-45, higher more active coping |
| **STAI** [SR 8,9] |  |  |  |  |
| Trait | Trait anxiety | 32.0±6.9 | 21-49 | 20-80 |
| State | State anxiety (pre experiment) | 31.5±6.2 | 22-51 | 20-80 |
| **MDMQ** [SR 10] |  |  |  |  |
| GoodBad A | Mood: Good vs bad (pre exp.) | 17.7±1.9 | 12-20 | 4-24, the higher the better |
| AwakeTired A | Mood: Awake vs tired (pre exp.) | 15.2±2.9 | 8-19 | 4-24, the higher the more awake |
| CalmNervous A | Mood: Calm vs nervous<br>(post exp.) | 16.5±2.6 | 10-20 | 4-24, the higher<br>the calmer |
| GoodBad B | Mood: Good vs bad (pre<br>exp.) | 17.2±1.7 | 12-19 | 4-24, the higher<br>the better |
| AwakeTired B | Mood: Awake vs tired<br>(pre exp.) | 10.1±2.4 | 6-16 | 4-24, the higher<br>the more awake |
| CalmNervous B | Mood: Calm vs nervous<br>(post exp.) | 17.2±2.1 | 12-20 | 4-24, the higher<br>the calmer |

**S2 Tab.** Average calibrated sound intensities 1 through 6, which were used as stimuli during the experiment. Cohort 2 received higher intensities, see Materials and Methods for rationale.

|  | Cohort 1<br>(n=15) |  |  | Cohort 2<br>(n=11) |  |  |
| --- | --- | --- | --- | --- | --- | --- |
| Intensity | Target VAS | dBA mean±SD | Range | Target VAS | dBA<br>mean±SD | Range |
| 1 | 25 | 73.3±5.4 | 61.0-82.9 | 48 | 85.0±9.0 | 65.7-99.3 |
| 2 | 35 | 77.0±4.9 | 67.7-85.9 | 59 | 91.3±8.3 | 71.1-101.5 |
| 3 | 45 | 80.5±5.0 | 73.6-89.6 | 70 | 95.1±7.6 | 76.3-102.5 |
| 4 | 55 | 83.8±5.4 | 76.7-96.1 | 82 | 97.6±6.1 | 81.7-103.0 |
| 5 | 65 | 86.6±5.4 | 78.9-99.3 | 93 | 99.3±4.8 | 86.3-103.0 |
| 6 | 75 | 89.1±5.2 | 81.0-100.5 | 105 | 100.6±3.5 | 91.0-103.0 |

**S3 Tab.** Differences of behavioral and psychological measures of the SCR-selected subsample compared to the not-selected subsample.

| Measure | Mean±SD of n=26<br>included subjects<sup>1</sup> | Mean±SD of n=32<br>excluded subjects | Mean differences<sup>2</sup> | Variance<br>differences<sup>3</sup> |
| --- | --- | --- | --- | --- |
| Heat pain threshold | 43.5±1.1 | 43.3±1.1 | t(56)=0.496,<br>p=0.622 | F(31,25)=1.000,<br>p=0.989 |
| Heat intensity 1 | 41.9±1.4 | 41.7±1.4 | t(56)=0.499,<br>p=0.620 | F(31,25)=1.094,<br>p=0.825 |
| Heat intensity 2 | 42.5±1.3 | 42.3±1.2 | t(56)=0.508,<br>p=0.613 | F(31,25)=1.066,<br>p=0.879 |
| Heat intensity 3 | 43.1±1.1 | 43.0±1.1 | t(56)=0.505,<br>p=0.615 | F(31,25)=1.024,<br>p=0.961 |
| Heat intensity 4 | 43.8±1.0 | 43.7±1.0 | t(56)=0.480,<br>p=0.633 | F(31,25)=0.975,<br>p=0.936 |
| Heat intensity 5 | 44.4±1.0 | 44.3±1.0 | t(56)=0.424,<br>p=0.673 | F(31,25)=0.930,<br>p=0.840 |
| Heat intensity 6 | 45.1±1.0 | 45.0±1.1 | t(56)=0.343,<br>p=0.733 | F(31,25)=0.905,<br>p=0.785 |
| Sound unpleasantness<br>threshold | 82.9±7.3 | 83.9±7.0 | t(56)=-0.537,<br>p=0.593 | F(31,25)=1.079,<br>p=0.853 |
| Sound intensity 1 | 76.8±10.4 | 78.3±9.2 | t(56)=-0.552,<br>p=0.583 | F(31,25)=1.283,<br>p=0.528 |
| Sound intensity 2 | 81.9±9.9 | 83.1±9.6 | t(56)=-0.454,<br>p=0.651 | F(31,25)=1.063,<br>p=0.885 |
| Sound intensity 3 | 86.2±9.8 | 86.7±9.5 | t(56)=-0.206,<br>p=0.838 | F(31,25)=1.055,<br>p=0.901 |
| Sound intensity 4 | 89.6±9.1 | 89.6±8.9 | t(56)=-0.004,<br>p=0.997 | F(31,25)=1.041,<br>p=0.928 |
| Sound intensity 5 | 92.4±8.0 | 92.0±8.1 | t(56)=0.172,<br>p=0.864 | F(31,25)=0.973,<br>p=0.932 |
| Sound intensity 6 | 94.7±6.9 | 94.0±7.3 | t(56)=0.369,<br>p=0.714 | F(31,25)=0.899,<br>p=0.771 |

<sup>1</sup>Included subjects pooled over both cohorts; <sup>2</sup>Two-sample t-test; <sup>3</sup>Two-sample F-test

**S1 Fig.**
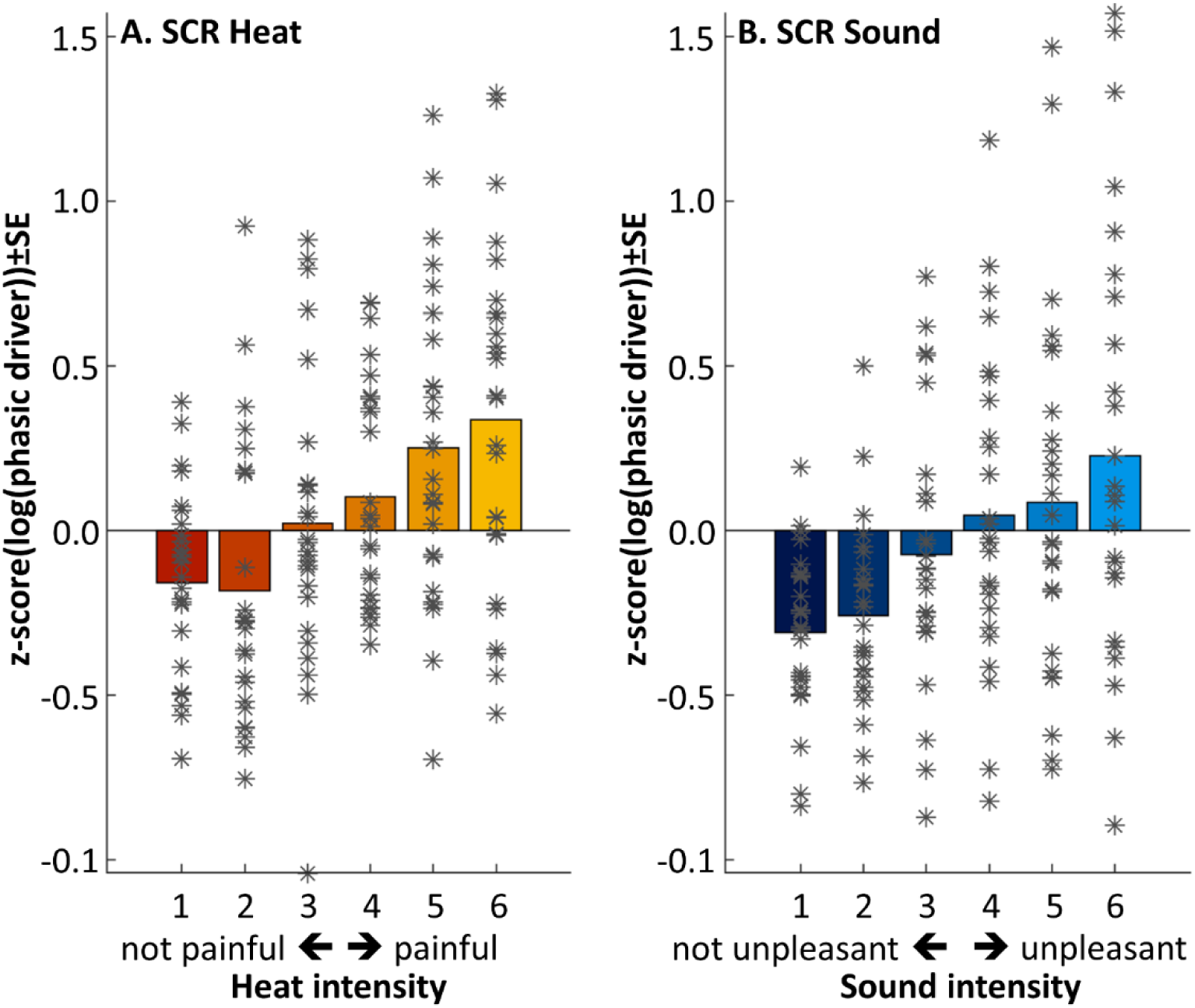
Skin conductance responses following heat (**A**) and sound stimuli (**B**), with individual data points. The pain and unpleasantness thresholds were located between intensities 3 and 4, as per calibration.

**S2 Fig.**
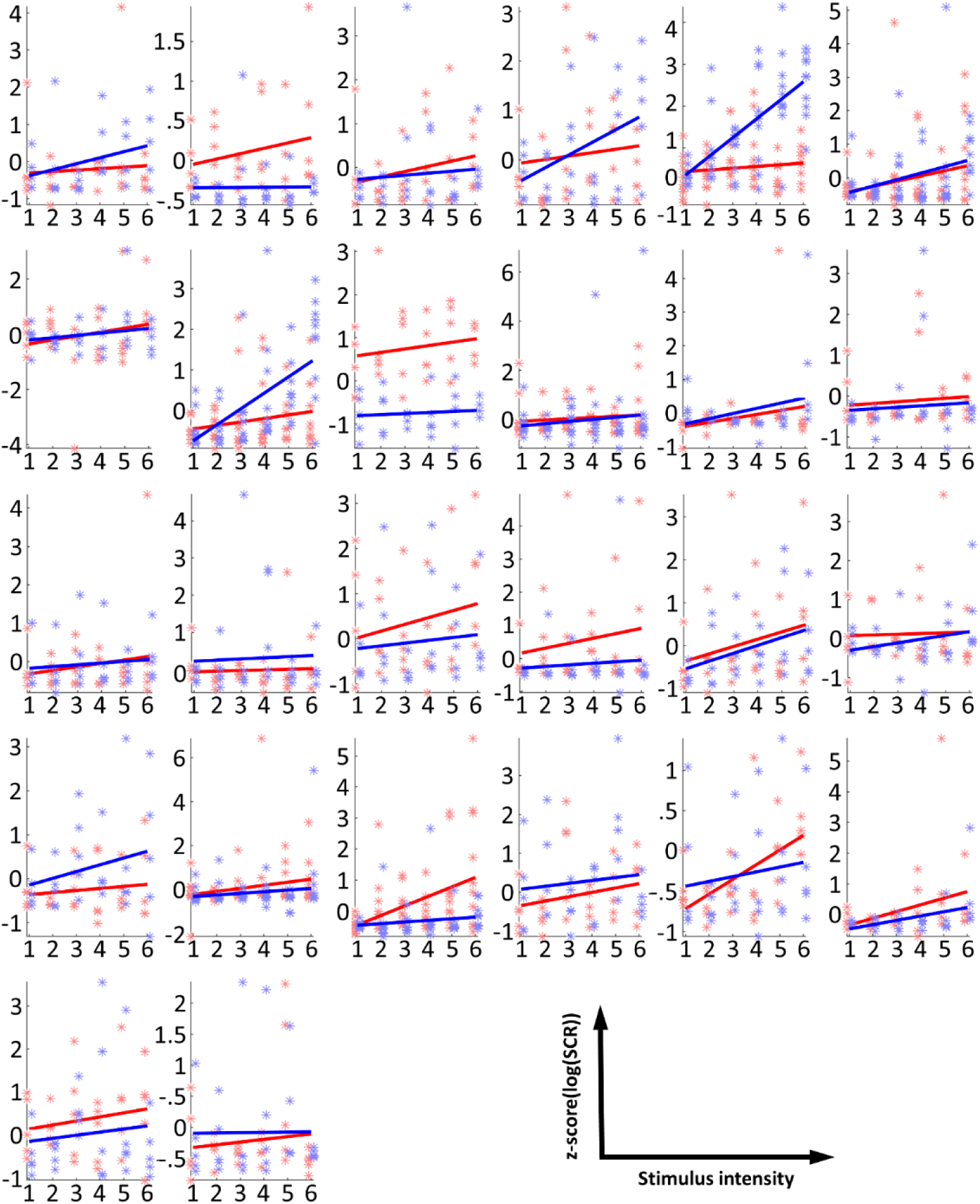
Single subject skin conductance responses following heat (red) and sound stimuli (blue). Asterisks indicate single responses, lines are regression curves as per linear regression. The pain and unpleasantness thresholds were located between intensities 3 and 4, as per calibration.

**S3 Fig.**
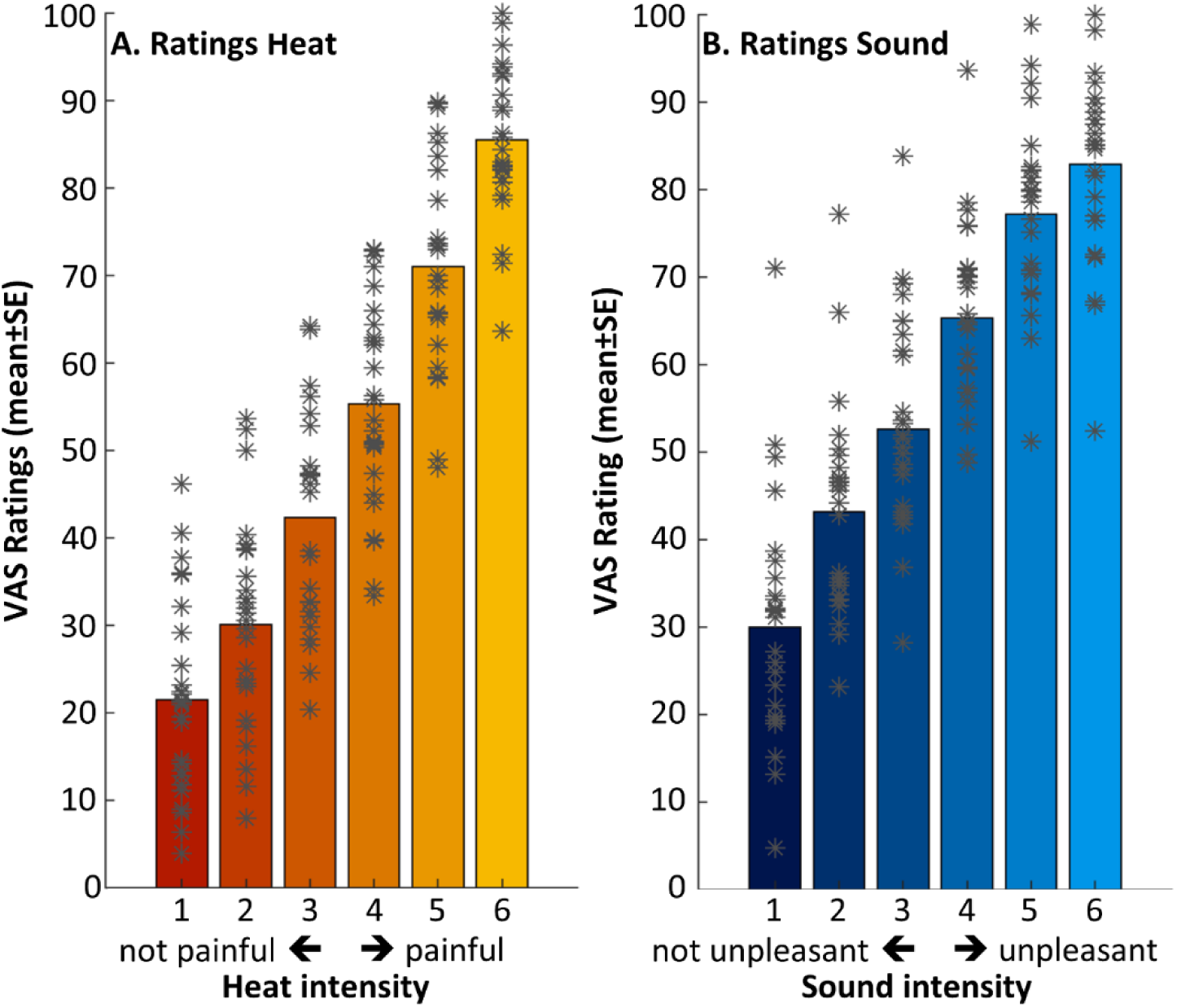
Behavioral ratings following heat (**A**) and sound stimuli (**B**), with individual data points. The pain and unpleasantness thresholds were located between intensities 3 and 4, as per calibration.

**S4 Fig.**
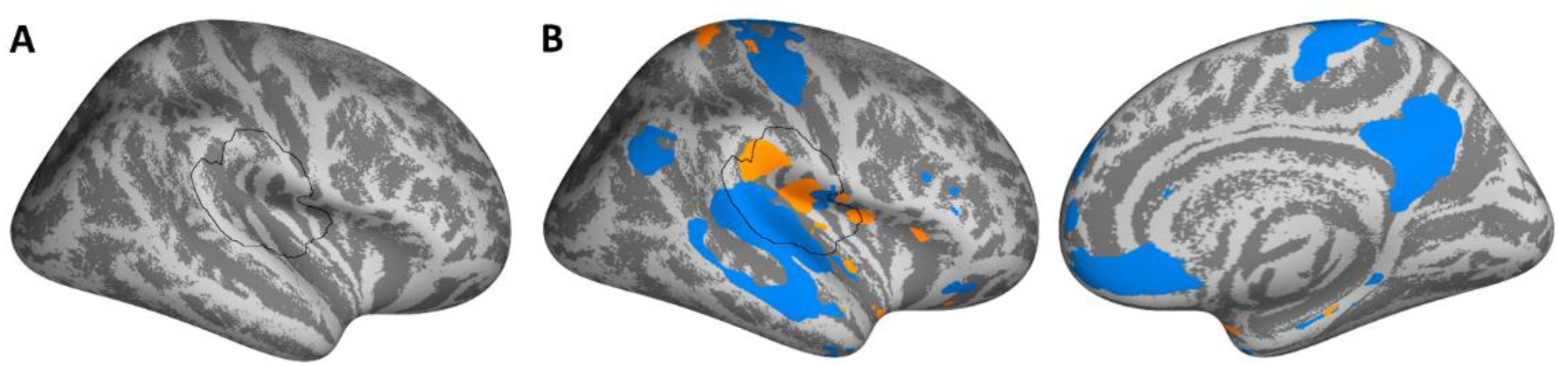
Binary and signed masks used for analyses. **A.** Binary mask used for small volume correction used for all analyses (unless otherwise noted), delineated by the black line. **B.** Signed mask used for covering heat (orange) or sound (blue) contrasts.

**S5 Fig.**
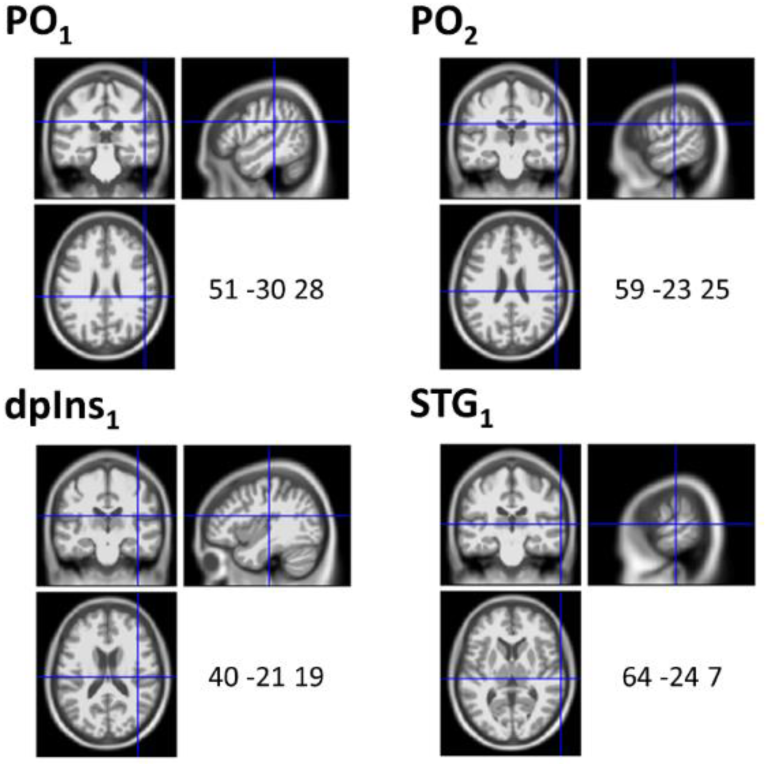
Location of peak voxels for modality main effects in the parietal operculum (PO1, PO2) and dorsal posterior insula (dpIns1) for heat, superior temporal gyrus (STG1) for sound. Also see Fig 4.

**S6 Fig.**
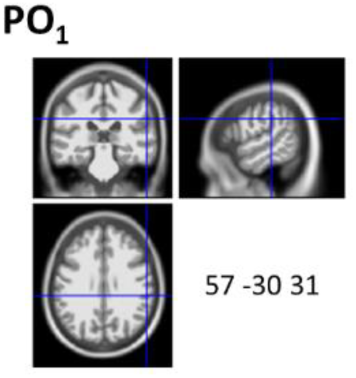
Location of peak voxel for parametric modulation by heat intensity in the parietal operculum (PO1). Also see Fig 5.

**S7 Fig.**
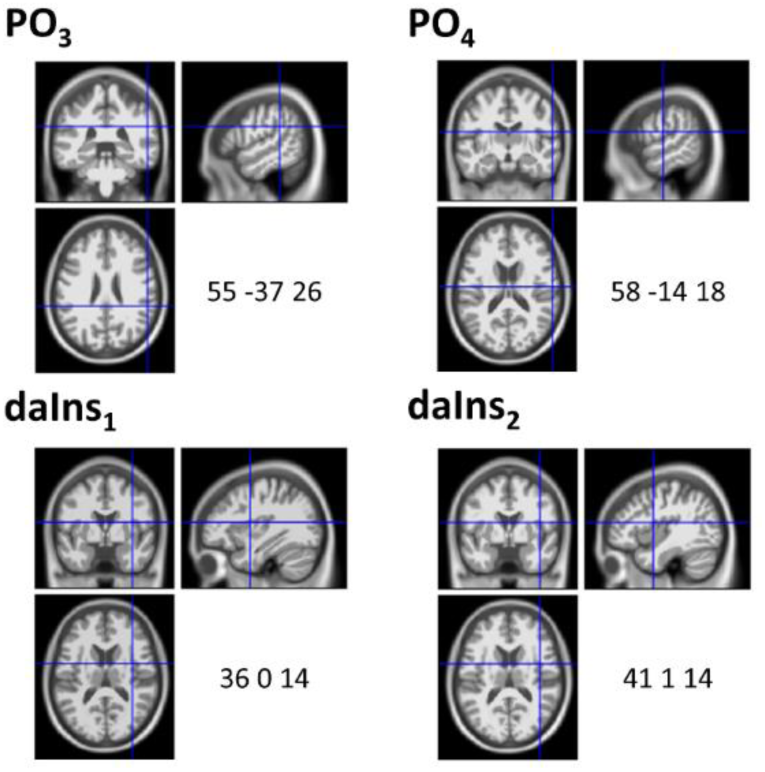
Location of peak voxels for parametric modulation by ratings in the parietal operculum (PO3, PO4) and dorsal anterior insula (daIns1, daIns2). Also see Fig 6.

**S8 Fig.**
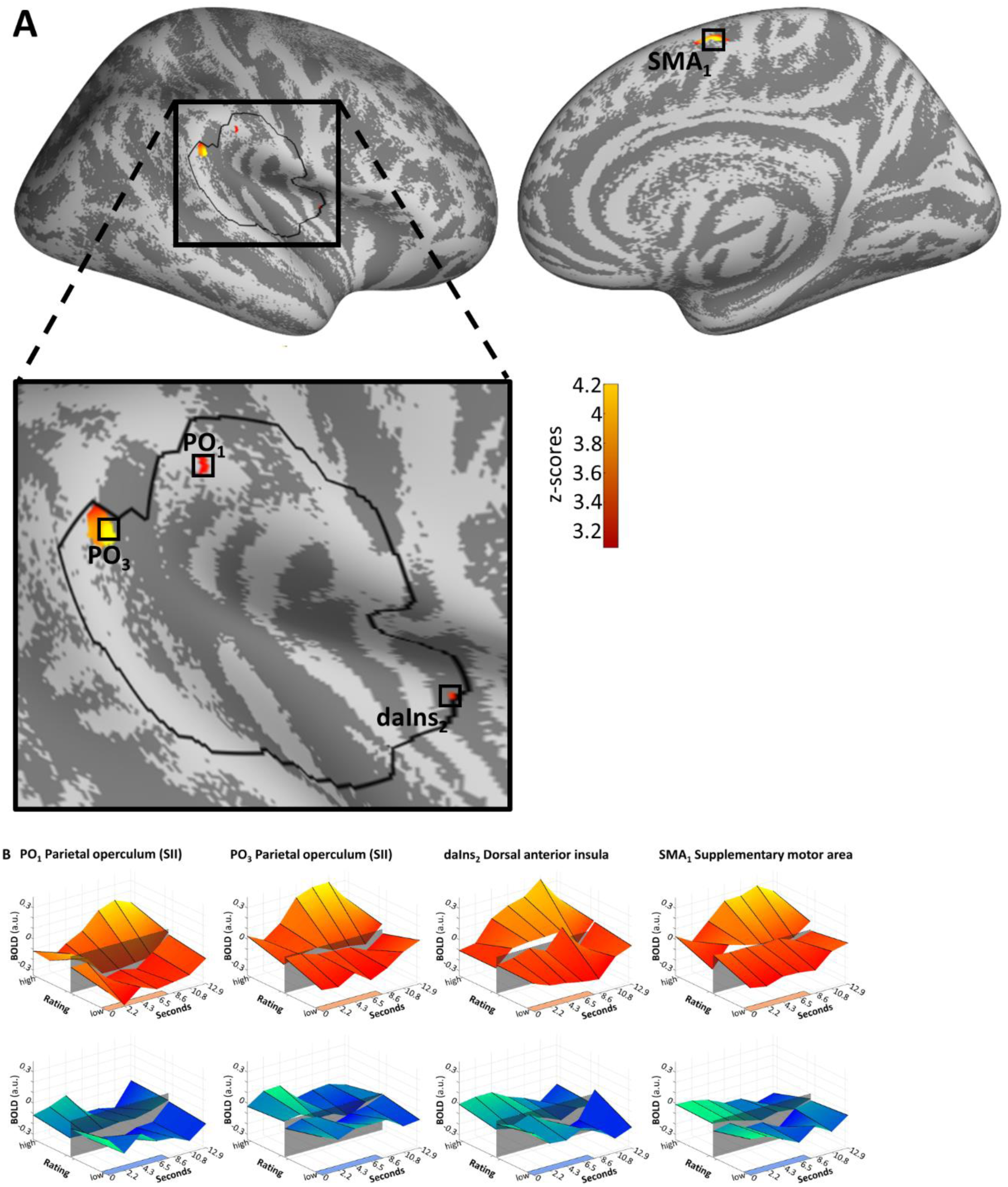
Medial surface view for areas that fulfill the axiomatic requirements of differential activation during pain compared to heat and sound (also see Fig 7). Unlike in Fig 7, activations are shown after analysis with a lower threshold of p(uncorrected)<0.001. At this threshold, activation was found in the parietal operculum (PO3 is the cluster reported in Fig 7; PO1 is a more dorsal cluster), the dorsal anterior insula (daIns2) and the (pre-)supplementary motor area (SMA1). Exact coordinates and activation of cluster peaks were x=55, y=-37, z=24, Z=4.176, p(uncorrected)=1e-05 (PO3), x=61, y=-26, z=30, Z=3.26, p(uncorrected)=6e-04 (PO1), x=35, y=1, z=13, Z=3.297, p(uncorrected)=5e-04 (daIns2), and x=7, y=-5, z=60, Z=4.202, p(uncorrected)=1e-05 (SMA1). **A**. Activations are thresholded at p(uncorrected)<0.001 and overlaid on an average brain surface. The black line delineates the SVC mask. **B**. Poststimulus plots of fMRI activation in the four vertices during heat (orange) and sound (blue). The shaded patch in the center signifies the pain threshold (for heat) and unpleasantness threshold (for sound). The colored patches at the right axes show the stimulus duration.

**S9 Fig.**
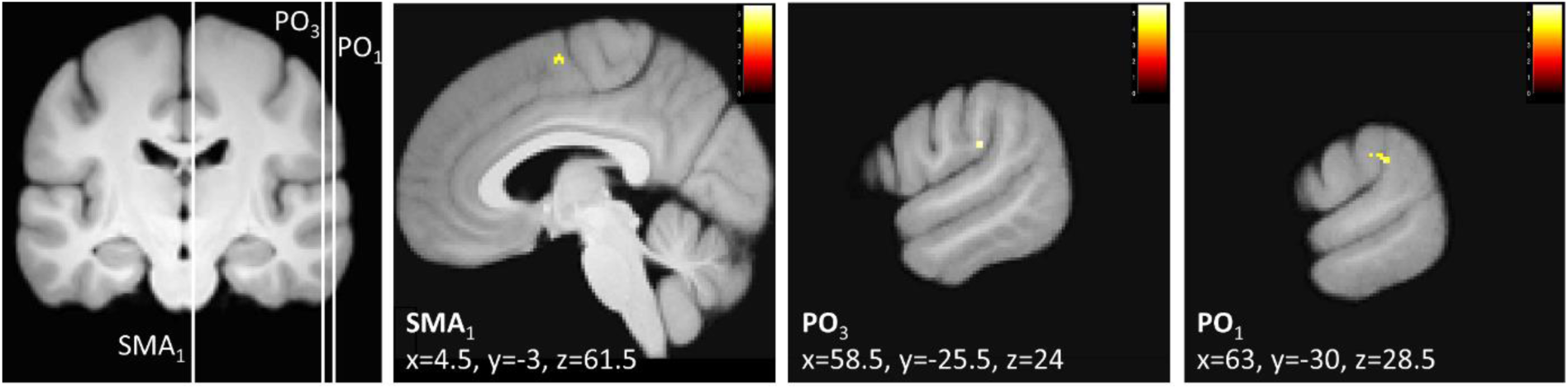
Conventional 3D analysis for areas that fulfill the axiomatic requirements of differential activation during pain compared to heat and sound (also see Fig 7). No activations were significant at our predefined threshold (p<0.05, corrected). For reasons of completeness, this analysis is reported at a lower threshold of p(uncorrected)<0.001. The location of activations roughly correspond to the surface analysis presented in S8 Fig. Activation was observed in the parietal operculum (PO3 is the cluster reported in Fig 7; PO1 is a more dorsal cluster) and the (pre-)supplementary motor area (SMA1), among other areas. Exact coordinates and activation of cluster peaks were x=59, y=-26, z=24, Z=3.9, p(uncorrected)=5e-05 (PO3), x=63, y=-30, z=29, Z=3.487, p(uncorrected)=2e-04 (PO1), and x=5, y=3, z=62, Z=3.297, p(uncorrected)=3e-04 (SMA1). Activations are thresholded at p(uncorrected)<0.001 and overlaid on an average brain.

**S10 Fig.**
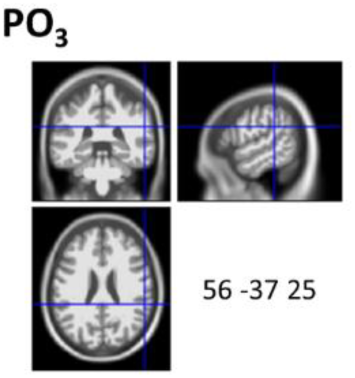
Location of peak voxel for activation corresponding to the four axioms in the parietal operculum (PO3). Also see Fig 7.

